# Diversity and Spatial Segregation of TRP Channels in Choanoflagellates Provide Insight into the Evolutionary Origin of Animal Sensory Systems

**DOI:** 10.64898/2026.06.15.732245

**Authors:** Simen Mannsåker, Pawel Burkhardt, Jeffrey Colgren

**Affiliations:** Michael Sars Centre, University of Bergen, Norway

## Abstract

Sensory systems, built around specialized cell types, are central to how animals perceive and respond to their environments. Yet many of the molecular components defining these systems predate the origin of animal multicellularity. Among these, transient receptor potential (TRP) channels form a polymodal and evolutionarily ancient superfamily of ion channels involved in diverse sensory processes. To better understand how sensory complexity emerged in animals, we investigated the diversity of TRP channels in choanoflagellates, the closest living relatives of animals. Using a combination of homology-based searches, phylogenetics, and structural predictions, we find extensive TRP channel repertoires across choanoflagellates, including representatives of most major animal TRP channel families. Comparative analyses across species revealed two distinct evolutionary patterns for TRP channel families: conserved, low-copy families with stable domain architectures, and lineage-specific expansions within the families TRPM and TRPW, indicative of functional diversification. Functional insights from fluorescent localization studies in the choanoflagellate *Salpingoeca rosetta* demonstrated that TRPA, TRPC, and TRPV channels are spatially segregated within the collar complex, a key interface for environmental sensing and feeding. Distinct localization domains, along with evidence for heteromeric interactions between TRPA paralogs, suggest that subcellular organization likely contributes to sensory specialization in these single cells. Together, our findings indicate that a diverse and functionally versatile TRP channel toolkit was already present in the last common ancestor of choanoflagellates and animals. We propose that the evolution of animal sensory systems involved both expansion and reorganization of this ancestral repertoire, with subcellular patterning in unicellular organisms representing a precursor to cell-type specialization in multicellular animals.

## Introduction

Animals rely on sophisticated sensory systems to detect, interpret, and respond to their environments. These systems enable external behaviors such as foraging, predator avoidance, navigation, and communication with conspecifics, while also monitoring internal physiological states. At the front end of these systems are often specialized sensory cells or neurons, which can be found across extant animal lineages (Arendt, 2021; Esposito, 2021; Schlosser, 2025). At the molecular level, sensory perception is mediated by receptor proteins that transduce physical and chemical cues into intracellular signals. Because sensory capabilities shape ecological niches and life-history strategies, sensory receptor families often undergo rapid diversification (Valencia-Montoya et al., 2024). At the same time, comparative genomics has revealed that many receptor superfamilies central to animal sensory biology originated before the emergence of animals (Valencia-Montoya et al., 2024), raising questions about their ancestral functions and how they were repurposed during the evolution of multicellularity.

Unicellular eukaryotes frequently exhibit complex environmental responsiveness, including taxis, mechanosensation, and chemosensation, despite lacking specialized sensory cell types (Ros-Rocher & Brunet, 2023; Wan & Jékely, 2021). In animals, sensory tasks are partitioned among differentiated cells and integrated through intercellular communication and electrical signaling. The transition to obligate multicellularity in animals therefore required not only the expansion of receptor repertoires, but also the reorganization of existing molecular components into new functional contexts (Colgren & Burkhardt, 2026). Identifying how these receptors are utilized in the close relatives of animals, and how they diversified within animal lineages, is central to understanding the evolutionary origins of animal sensory systems.

Choanoflagellates are the sister group to animals, and therefore, essential to understanding the evolutionary events that led to the transition to animal multicellularity (Brunet & King, 2017; Carr et al., 2008; Nitsche et al., 2011). With more than 360 described species, choanoflagellates comprise a morphologically and ecologically diverse clade of bacterivorous protists characterized by a polarized cell body bearing a single apical flagellum surrounded by a collar of actin-based microvilli (Leadbeater, 2015; Richter & Nitsche, 2017). Beating of the flagellum generates water currents that concentrate bacteria at the collar, where they are captured and phagocytosed (**Figure 1a**) (Dayel & King, 2014; Leadbeater, 2015). Choanoflagellates are cosmopolitan and found in a diverse range of aquatic habitats, can be swimming or benthic, and can have organic or silica-based extracellular layer (Leadbeater, 2015; Richter & Nitsche, 2017). Furthermore, they can have complex lifecycles, including states of clonal multicellularity (Dayel et al., 2011). The microscopic environments that choanoflagellates navigate are highly heterogenous, and the cells have been shown to respond to many environmental parameters, including pH, oxygen content, physical confinement, light, flow, and several molecules produced by bacteria (Brunet et al., 2021; Brunet et al., 2019; Dayel et al., 2011; Ireland Ella et al., 2020; Kirkegaard et al., 2016; Miño et al., 2017; Ros-Rocher & Brunet, 2023; Stocker, 2012; Woznica et al., 2016; Woznica et al., 2017). Additionally, electrical signaling has recently been demonstrated to be associated with feeding in the choanoflagellate model *Salpingoeca rosetta,* and was found to function in coordinating behavior in multicellular rosettes (Colgren & Burkhardt, 2025). Furthermore, genomic and transcriptomic data have revealed a rich repertoire of membrane receptors, including many G protein-coupled receptors and receptor tyrosine kinases (Garcia De Las Bayonas & King, 2025; King et al., 2008).

**Figure 1:**
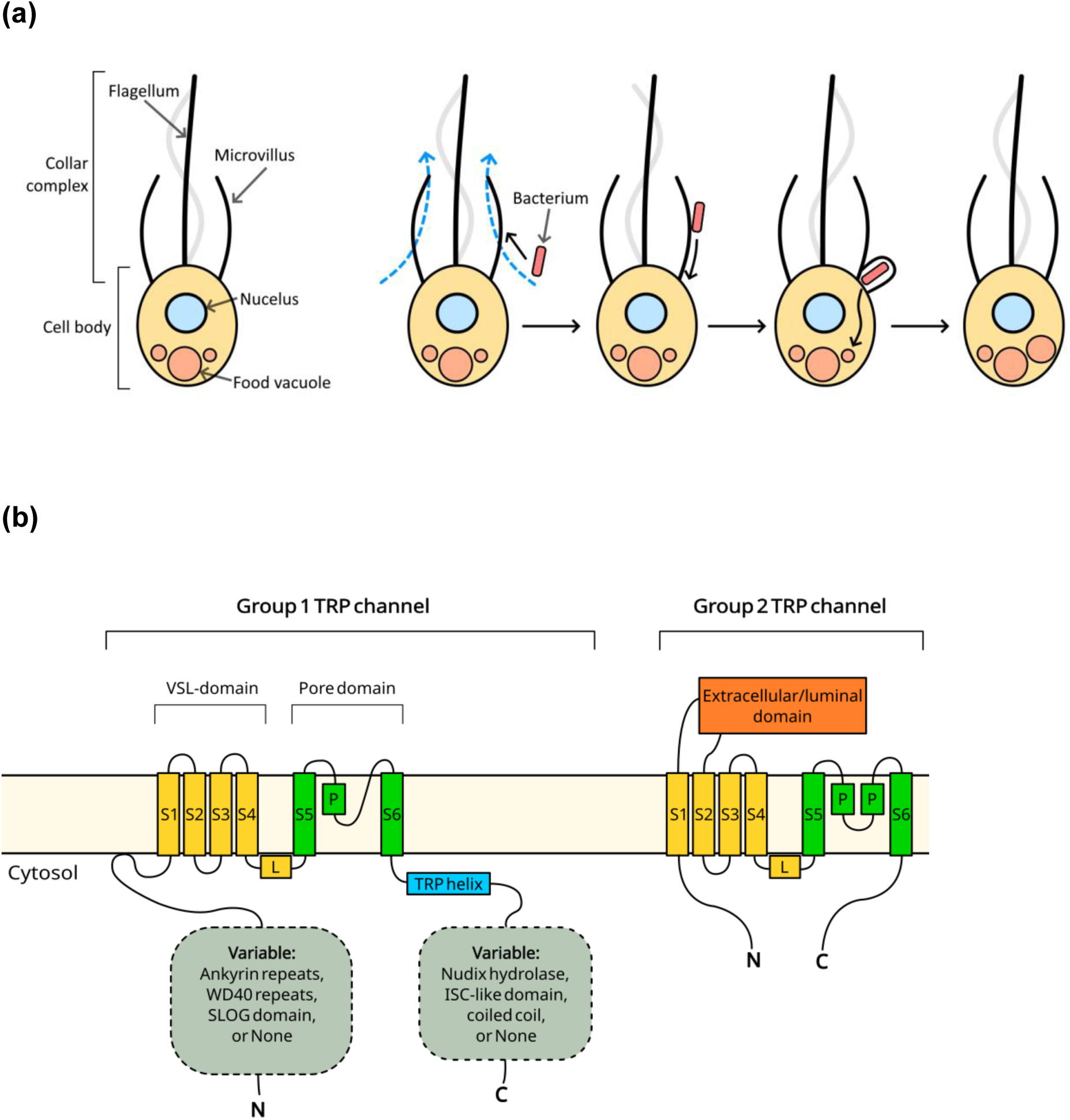
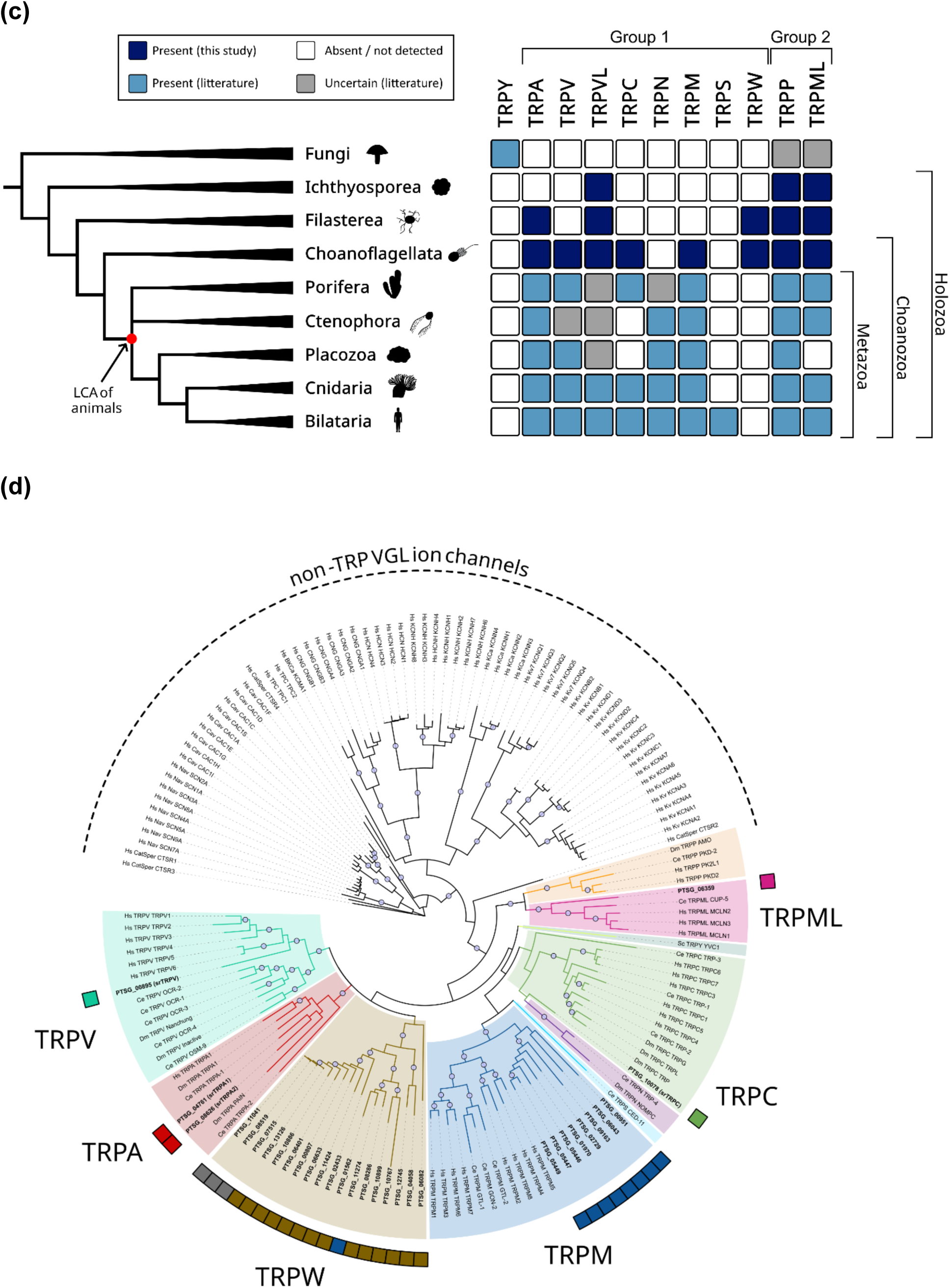
Framework for TRP channel diversity and collar complex in *S. rosetta*. **(a)** Choanoflagellates use the collar complex, consisting of a flagellum and a ring of microvilli (only two microvilli depicted) called the collar, for feeding. Flagellar beating creates water currents (blue) that draw in bacteria (red) into the distal part of the collar. Captured particles are transported to the base of the collar and then phagocytosed. The bacteria are digested in basal food vacuoles. **(b)** A simplified topology for domain organization of TRP channel monomers. The monomers assemble into tetramers with a central ion pore. TRP channels have diverse extramembrane compositions but share a common six-transmembrane helix (S1-S6) topology. The TMD is separated by a linker helix (L) into a voltage-sensing-like (VSL) domain, and the pore domain. Group 2 TRP channels have extracellular/luminal domains, and usually a split pore helix (P). Group 1 TRP channels usually have a TRP helix and elaborate cytosolic domains, with strong variation both between and within each TRP channel family. Most TRP channels do not have large domains in the C-terminal region, but some have enzymatic domains like Nudix hydrolase domains or isochorismatase-like (ISC-like) domains. **(c)** A cladogram of major holozoan lineages, showing the distribution of various TRP channel families. For each lineage, the families were marked as either present according to this study (dark blue), present according to literature search (light blue), absent (or not yet reported) (white), or uncertain (grey) if literature search yielded conflicting results (Davila-Velderrain & van Giesen, 2025; Himmel & Cox, 2020; Himmel et al., 2020; Hsiao et al., 2021; Jaślan et al., 2020; Peng et al., 2015). **(d)** A TMD-based maximum-likelihood phylogenetic tree of putative *S. rosetta* TRP channels in conjunction with TRP channels and non-TRP voltage-gated-like (VGL) ion channels from other organisms (*Saccharomyces cerevisiae* (Sc), *H. sapiens* (Hs), *C. elegans* (Ce) and *D. melanogaster* (Dm)). The branches of the TRP channel clades are colored based on family inferred from phylogeny. Peripheral boxes mark *S. rosetta* entries, and the coloring refers to the expected classification based on domain predictions and a reciprocal BLASTP search. For the TRPW group, entries with brown coloration had WD40-repeat-containing proteins as top annotated hits in BLAST search, while candidates without known extramembrane domains were marked with grey filling. Bootstrapping values (UFBoot) were indicated as circles along the branch if greater than 0.95.

Transient receptor potential (TRP) channels form a large and diverse superfamily of cation channels with well-established roles in animal sensory physiology, including the detection of temperature, mechanical stimuli, osmolarity, and a wide range of chemical ligands (Julius, 2013; Zhang et al., 2023). The TRP channel superfamily is evolutionarily ancient and present across most major eukaryotic lineages, with the exception of plants (Himmel & Cox, 2020). Cation selectivity, extramembrane domain composition and the factors regulating gating are all highly variable within the superfamily. A unifying feature is their polymodal nature, meaning that gating is affected by a multitude of factors, including physical stimuli and numerous exogenous and endogenous compounds (Abe & Puertollano, 2011; Bellemer, 2015; Cox et al., 2024; Julius, 2013; Kashio & Tominaga, 2022; Laursen et al., 2015; Lu et al., 2022; Startek et al., 2019; Talyzina et al., 2024; Zhang et al., 2023; Zheng, 2013). Consequently, TRP channels act both as primary transducers of sensory stimuli and as integrators of pathways initiated by other proteins, acting as a flexible evolutionary framework for sensory responsiveness.

Structurally, TRP channels are related to voltage-gated ion channels and assemble as tetramers, with each subunit generally containing six transmembrane helices and a pore-forming loop between helices S5 and S6 (**Figure 1b**). Large cytosolic or luminal/extracellular domains, which differ markedly among families, contribute to channel assembly, ligand binding, and allosteric regulation (Liao et al., 2013; Schindl & Romanin, 2007; Won et al., 2025; Zhang et al., 2023; Zheng, 2013). Phylogenetic analyses indicate an ancient divergence between the two groups of extant TRP channels (Himmel & Cox, 2020). In animals, these can be distinguished by whether extramembrane domains are cytosolic or luminal. Animal TRP channels have traditionally been classified into seven families: TRPC, TRPA, TRPM, TRPV and TRPN belonging to Group 1, and TRPML and TRPP belonging to Group 2. The Group 1 families TRPVL and TRPS have been recognized more recently (Himmel et al., 2020; Peng et al., 2015; Schüler et al., 2015; Zhang et al., 2023). Most non-animal TRP channels do not clearly fit these families, although previous studies have described the presence of animal-like TRP channels in choanoflagellates (**Figure 1c**) (Ahmed et al., 2022; Cai, 2008; Davila-Velderrain & van Giesen, 2025; Iordanov et al., 2019; Lindström et al., 2017; Peng et al., 2015; Sigg et al., 2017). However, with the exception of *in vitro* characterization of a TRPM member (Huang et al., 2024; Iordanov et al., 2019), the previous investigations are largely limited to annotation based on similarity to human orthologs, and have been confined to only two species of choanoflagellates.

Here, we systematically analyze TRP channel repertoires across choanoflagellates to reconstruct the evolutionary history and TRP channel family diversity likely present in the unicellular ancestor of animals. Furthermore, we examine the subcellular localization and interactions of orthologs of the holozoan-specific families TRPA, TRPC, and TRPV in *S. rosetta*. We find all channels localized to the flagellar-collar apparatus, but with discrete boundaries between the different families and evidence for heteromeric interaction between two TRPA members in the flagella. This study advances our understanding of the TRP channel diversity present in the last common ancestor of choanoflagellates and animals, and provides new hypotheses on how the subsequent diversification has shaped extent sensory systems. More broadly, it offers insights into how ancient, polymodal ion channels were co-opted and elaborated during the emergence of animal multicellularity and sensory complexity.

## Results

### Characterization of the TRP channels in *S. rosetta*

Focusing on putative transmembrane domains (TMDs), we identified 31 candidate TRP channels in the *S. rosetta* genome (Fairclough et al., 2013), which grouped with animal TRP channels (**Figure 1d**). Among these were one member each of TRPML, TRPV, and TRPC, and two members of TRPA families (hereafter called *sr*TRPML (PTSG_06359), *sr*TRPV (PTSG_00895), *sr*TRPC (PTSG_10078), *sr*TRPA1 (PTSG_04761) and *sr*TRPA2 (PTSG_08626)). Full domain predictions showed consistency with animal orthologs, with the exception of an isochorismatase-like domain near the C-terminus of *sr*TRPV. Homotetramers were modeled (Abramson et al., 2024) for these five putative channel proteins, indicating expected assembly and a central ion pore (**Figures S1-S5**). Features such as a TRP helix, ankyrin repeats and coiled coil motifs could also be clearly identified from these predicted structures. The highest sequence similarity with animal orthologs was generally found in the pore-forming helices, and structural alignment of the predicted TMDs with human orthologs showed a very high degree of structural similarity (**Figure S6**). Together, these observations support a conserved role as ion channels. Note that the predicted sequence for *sr*TRPA2 (F2UK79_SALR5) lacked the S4-S5 linker region, which would disrupt the TMD. However, targeted sequencing of cDNA extracted from our cultures showed that this was due to an 83-residue region lacking from the genomic sequence (**Figure S7**).

Eight of the *S. rosetta* hits were classified as TRPMs based on the phylogenetic placement. Most were predicted to retain a canonical cytosolic N-terminal organization, including a SLOG domain, although one protein (PTSG_01970) was predicted to have a truncated N-terminal region. Additional variation was observed among individual members, with two sequences (PTSG_00951 and PTSG_09163) showing increased sequence divergence, and PTSG_00951 predicted to contain many additional short regions not found in other TRPMs. A C-terminal NUDIX hydrolase domain, characteristic of human TRPM2, was identified in five of the TRPMs. Together, these observations indicate an expanded but structurally heterogeneous TRPM repertoire in *S. rosetta*.

The remaining 18 putative *S. rosetta* TRP channels formed a monophyletic branch within the Group 1 TRP channels (**Figure 1d**). Though there was considerable variation outside the TMD of these proteins, 14 of them contained one or more cytosolic β-propeller WD40 repeat domains in their N-terminal region. Although AlphaFold predictions did not generate high-confidence models, the canonical propeller structure was present. WD40 repeat domains are commonly associated with protein-protein interactions, though not typically found in ion channels. This clade was tentatively named TRPW.

Of the four proteins in this clade that lacked WD40 repeat domains, one (PTSG_11274) contained both the NUDIX hydrolase and SLOG domain common in TRPMs, as well as three additional TM helices near the N-terminus. The three remaining proteins were highly similar in sequence to each other, lacked detectable domains outside of the TMD, and two of them lacked the helices S1-S2. These divergent proteins, combined with high variability among those with WD40 repeat domains, suggest a high structural diversity within the TRPW clade.

### Diverse TRP Channel Repertoires and Expansions within Choanoflagellates

To better characterize the diversity of TRP channels in choanoflagellates a custom TMD-focused HMM was used to search for orthologs in transcriptomic and genomic data derived from 20 additional species (Brunet et al., 2019; King et al., 2008; Richter, 2018). The HMM retrieved 393 entries, of which 330 remained after filtering based on predicted transmembrane topology to remove highly truncated sequences and members of closely related superfamilies with differing topology. These were then assigned to families based on sequence similarity network analysis and phylogenetic placement with well-annotated animal TRP channels (**Figure 2a, S8**). All families found in *S. rosetta* were represented in other species, with TRPM and TRPW showing high abundance. Contrary to previous studies (Cai, 2008; Davila-Velderrain & van Giesen, 2025; Peng et al., 2015), we also found putative TRPPs and TRPVLs in choanoflagellates. Intriguingly, the TRPPs appear to be the family with the highest degree of similarity between animal and choanoflagellate orthologs, with sequence identities with human TRPPs reaching 40%. Outside of the TMD, domain predictions revealed conserved architectures in most families, while some divergence could be found in TRPMs and TRPWs (**Figure 2b**). However, while TRPML transmembrane regions appear well-conserved amongst choanoflagellates, only half of the identified proteins had a predicted ion transport domain. Among the potentially enzymatic domains, the NUDIX hydrolase domain was found in one fifth of TRPMs. The isochorismatase-like domain was present in 14 of 18 Craspedida TRPVs, while absent from the Acanthoecida proteins (**Figure 2b)**.

**Figure 2:**
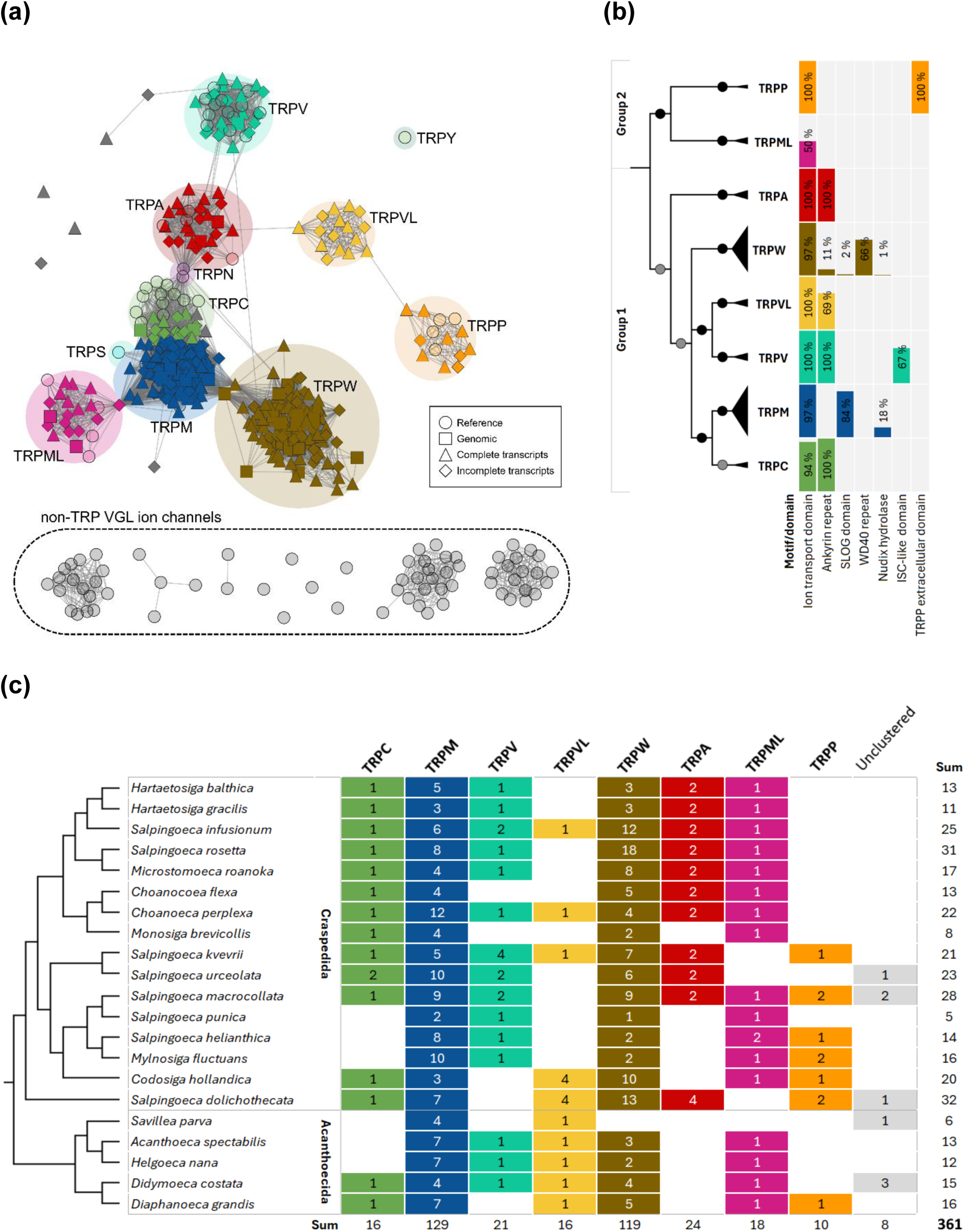
Diversity of TRP channels among choanoflagellates. **(a)** A sequence similarity network (SSN) of full-length peptide sequences including 349 peptide sequences from 21 choanoflagellate species. Well-annotated TRP channels from *H. sapiens*, *D. melanogaster*, *C. elegans* and *S. cerevisiae* and non-TRP voltage-gated-like (VGL) ion channels from *H. sapiens* were also included for reference (nodes shown as circles with opaque filling). The choanoflagellate entries were derived from genomic sequences (square nodes), or complete (diamond) or incomplete (triangle) transcriptomic sequences. Edges connect entries with pairwise BLASTP E-values less than 10^-20^. Clustering in the SSN strongly matches the placement in a TMD-based phylogenetic tree (node color based on phylogenetic placement in **Figure S8**). None of the hits clustered with the reference non-TRP VGL ion channels. The TRPVL cluster did not include any references, but a BLAST search showed strong similarity to TRPVL in *Nematostella vectensis*. **(b)** A cladogram derived from the TMD-based maximum-likelihood phylogenetic tree (**Figure S8**) showed strong association between family grouping and domain content. The phylogenetic tree suggested that all but eight putative choanoflagellate TRP channels could be placed in one of eight discrete branches with high Bootstrap support (UFBoot: black circle: ≥95%; grey circle: ≥80%). The Pfam HMMs were used to examine domain composition for each family and bar heights indicate the percentage of entries with each domain or motif. For the TRPP extracellular domain, a custom HMM built using animal homologs was applied. The ISC-like domain refers to an isochorismatase-like domain. **(c)** The distribution of each family is shown across the 21 choanoflagellate species. The evolutionary relationship between species is depicted as a cladogram derived from (Carr et al., 2017). Results for *S. rosetta* and *M. brevicollis* were inferred from genome-derived proteomes, while transcriptomic data were used for all other species.

The putative TRP channel repertoire was variable between the various choanoflagellate species, with no single species containing the full complement of identified families (**Figure 2c**). This is likely at least partly due to primary screening being performed on transcriptomes, thus missing lowly or non-expressed genes. The only family found in every species analyzed was TRPM. Apart from presence and absence, there was also variability in the number of proteins in each family between the species. When present, the TRPC, TRPV, TRPVL, TRPA, TRPML, and TRPP families generally had one or two members per species, and never more than 4 (**Figure 2c**). Interestingly, TRPA members were always found in pairs when present (or 4 members in the case of *Salpingoeca dolichothecata*). We find two choanoflagellate clades within the TRPA family (referred to as TRPA1 and TRPA2), which are nearly evenly represented (**Figure S9**). For all families except TRPP, the animal reference TRP channels form early-branching monophyletic groups within each family, suggesting that radiations in choanoflagellates and animals have occurred independently.

Alignments of choanoflagellate members within TRP channel families revealed varied levels of residue conservation (**Figures S10-S11**). The pore-forming helices, the TRP helix and cytosolic ankyrin repeats were generally well-conserved, albeit with some more divergence regions for the latter. These patterns would be consistent with maintaining ion permeability and certain interaction partners. In contrast, the voltage-like sensing domain of TRPC and TRPV had more variable sequences, suggesting looser evolutionary constraints. When present, the core of the isochorismatase-like domain of TRPVs also showed little variation. The characteristic Group 2 luminal domain of TRPML was made up of a mixture of well-conserved and variable residues, and the cytosolic regions showed large interspecific variation in length and sequence.

### Divergent TRP Channel Repertoires across Holozoan Lineages

To place the TRP channels of choanoflagellates in a broader evolutionary context, a search for orthologs was also done in genome-derived proteomes of ten other non-animal holozoans (**Figure 3a**). These included ichthyosporeans and filastereans, both having key phylogenetic positions for understanding the origin of animals (Brunet & King, 2017; Sebé-Pedrós et al., 2013; Shabardina et al., 2024).

**Figure 3:**
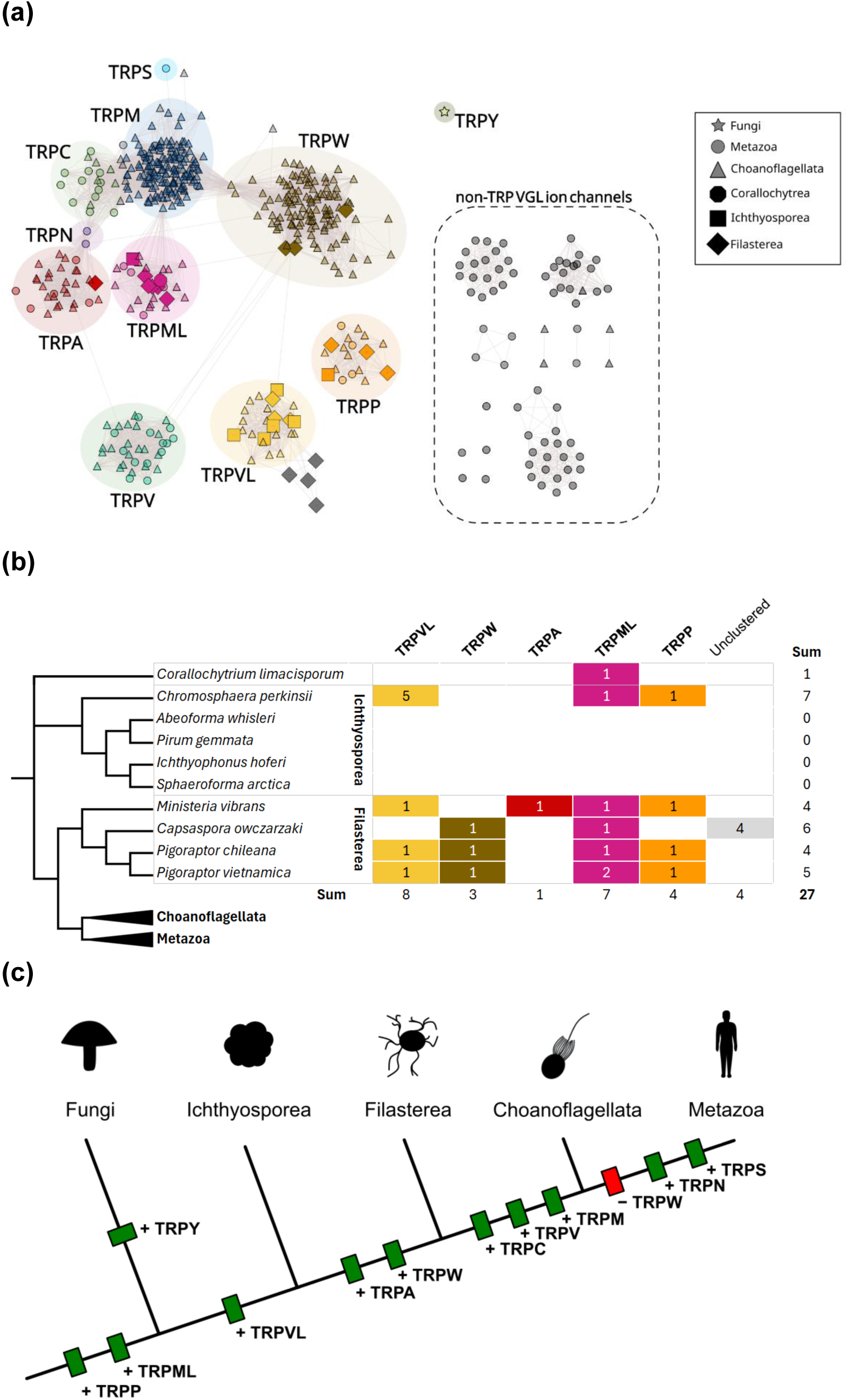
TRP channels in non-choanozoan holozoans. **(a)** A sequence similarity network (SSN) of full-length peptide sequences including all identified sequences from Corallochytrea, Ichthyosporea and Filasterea. The sequences were identified from genome-derived protein sequences through a TMD-focused HMM. Animal and choanoflagellate TRP channels, and non-TRP voltage-gated-like (VGL) ion channels from *H. sapiens* were included for reference. Coloring of nodes in the SSN was based on phylogenetic placement in a TMD-based phylogenetic tree (**Supplemental file S6**). None of the hits clustered with the non-TRP VGL ion channels. **(b)** The distribution of each detected family is shown for 10 non-choanozoan holozoan species. The evolutionary relationship between species is depicted as a cladogram derived from (Liu et al., 2024; Shabardina et al., 2024). **(c)** A cladogram of opisthokonts (holozoans and fungi) and a scheme of appearance and clade-wide loss of TRP channel families that would be consistent with the results described in the main text. Note that it is currently unclear whether TRPP-like and TRPML-like proteins in fungi are true TRPPs and TRPMLs. Note also that our results do not rule out the possibility of earlier origin of the TRP channel families due to limited sampling.

For four of the five ichthyosporean genomes explored, no TRP channels could be found. *Chromosphaera perkinsii*, however, possessed several animal-like TRP channels, including one TRPP, one TRPML and five TRPVLs (**Figure 3b**). Notably, this species also differs by being free-living and belonging to the early-branching clade Dermocystida, while the other four species are parasitic ichthyophonids (Glockling et al., 2013; Grau-Bové et al., 2017). The lack of TRP channels in the examined ichthyophonids likely reflects losses, although it remains uncertain whether this is clade-wide, or has occurred several times within the clade. The absence could potentially relate to a lifestyle with relaxed selective pressures for broad environmental sensing and signal integration due to living within a host.

The four species of filastereans examined revealed the presence of TRPML, TRPP, TRPA, TRPW and TRPVL, albeit not in all species. TRPW family members are present in three of the species, but only with one sequence per species **(Figure 3b**). Sequence comparisons of filasterean TRPW members show close similarity, contrasting the divergence and radiation of TRPWs observed within Choanoflagellata. Each of these contained predicted WD40 repeat-containing domain, suggesting that TRPWs are not unique to choanoflagellates, and were likely lost in the animal lineage (**Figure 3c**).

### Spatial segregation of TRP channels within the collar complex of *S. rosetta*

Choanoflagellates are highly polarized cells which maintain stable functional regionalization, most prominently at the apical flagellar and collar domains. In order to gain further insights into the orthologs of animal TRP channels in choanoflagellates, we examined their subcellular localization. The *S. rosetta* members of TRPA, TRPC and TRPV (*sr*TRPA1, *sr*TRPA2, *sr*TRPC and *sr*TRPV) were examined *in vivo* via fluorescent overexpression. These three families were chosen due to a strong relevance to animal sensory neurons, while also having a limited number of paralogs in *S. rosetta*. Each protein was cloned into an *S. rosetta* expression construct with the fluorescent protein StayGold attached to the C-terminus (Karapidaki et al., 2026).

After transfection and recovery (Booth et al., 2018), *S. rosetta* cultures were plated and live imaged to examine protein localization. This revealed distinct subcellular localization patterns for each of the four TRP channel fusion proteins (**Figure 4**). *sr*TRPA1 was the only one that seemed to be restricted to internal membranes, and the signal was particularly strong in the ER. This could point to a role in intracellular calcium homeostasis or result from improper trafficking and accumulation when highly expressed. The closely related *sr*TRPA2, on the other hand, displayed a strong signal along the flagellum. *sr*TRPC localized to the microvillar collar, while showing no detectable signal in the flagellum. The signal was particularly strong at the distal parts of the microvilli, while not detectable at the base. *sr*TRPV also showed localization to the collar region, which was strongest at the base and the membrane surrounding the collar. The signal weakened and becomes undetectable towards the tip of the microvilli. Thus, *sr*TRPA2, *sr*TRPC, and *sr*TRPV all localized to the collar complex, while showing little overlap within the structure.

**Figure 4:**
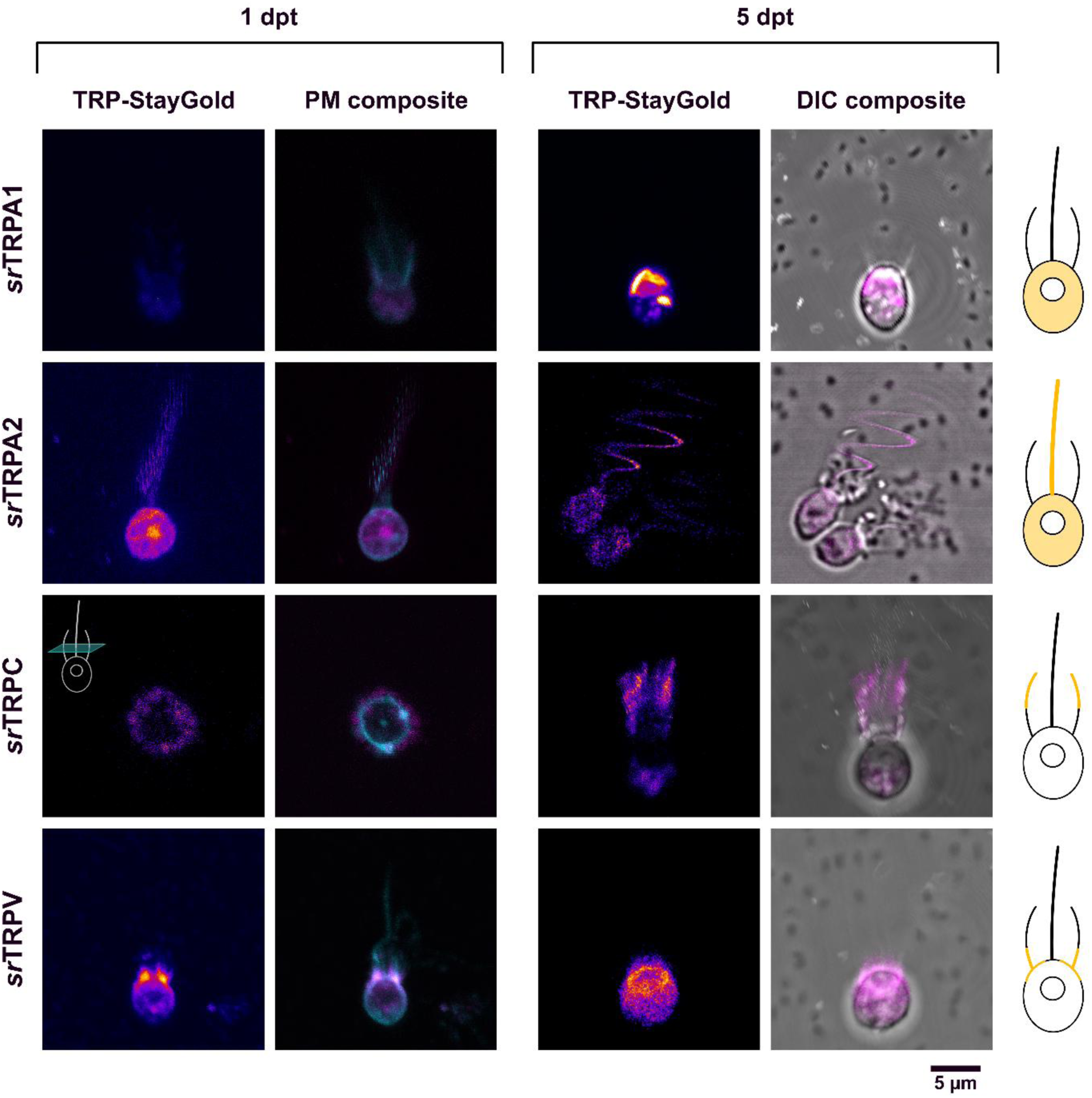
Distinct localization patterns of fluorescently tagged TRP channels in *S. rosetta*. The cells were transfected with a plasmid encoding an endogenous TRP channel fused to the fluorophore StayGold and imaged live by confocal microscopy either 1 or 5 days post-transfection (dpt). To place the TRP-StayGold signal (magenta in composites, fire LUT when shown alone) in a cellular context, fluorescence signal was overlaid either signal from a plasma membrane-tagging fluorophore (cyan in PM composite), or a differential interference image (DIC composite). The observed localization patterns for each TRP channel were summarized in the cartoon schematics. The cellular orientation of the 1 dpt image for *sr*TRPC is orthogonal to the other cells, as depicted by inset schematic, to better visualize the collar signal.

To further investigate the localization patterns of these TRP channels, we generated mCherry fusions of each and co-expressed pairs of different channels. We observed localizations consistent with the single overexpression for the combinations of *sr*TRPC and *sr*TRPV, as well as for *sr*TRPC and *sr*TRPV with either of the TRPA paralogs. However, when *sr*TRPA1 was co-expressed with *sr*TRPA2, there was an accumulation of the signal in the flagellum, which was not clearly visible when *sr*TRPA1 was over-expressed alone (**Figure 5**). This occurred regardless of which fluorophore was fused to which channel. In these cases, *sr*TRPA2 showed a consistent localization in the cell body and along the flagellum as observed when over-expressed alone. This could point to an interaction between the two paralogs, potentially including the formation of heteromeric channels.

**Figure 5:**
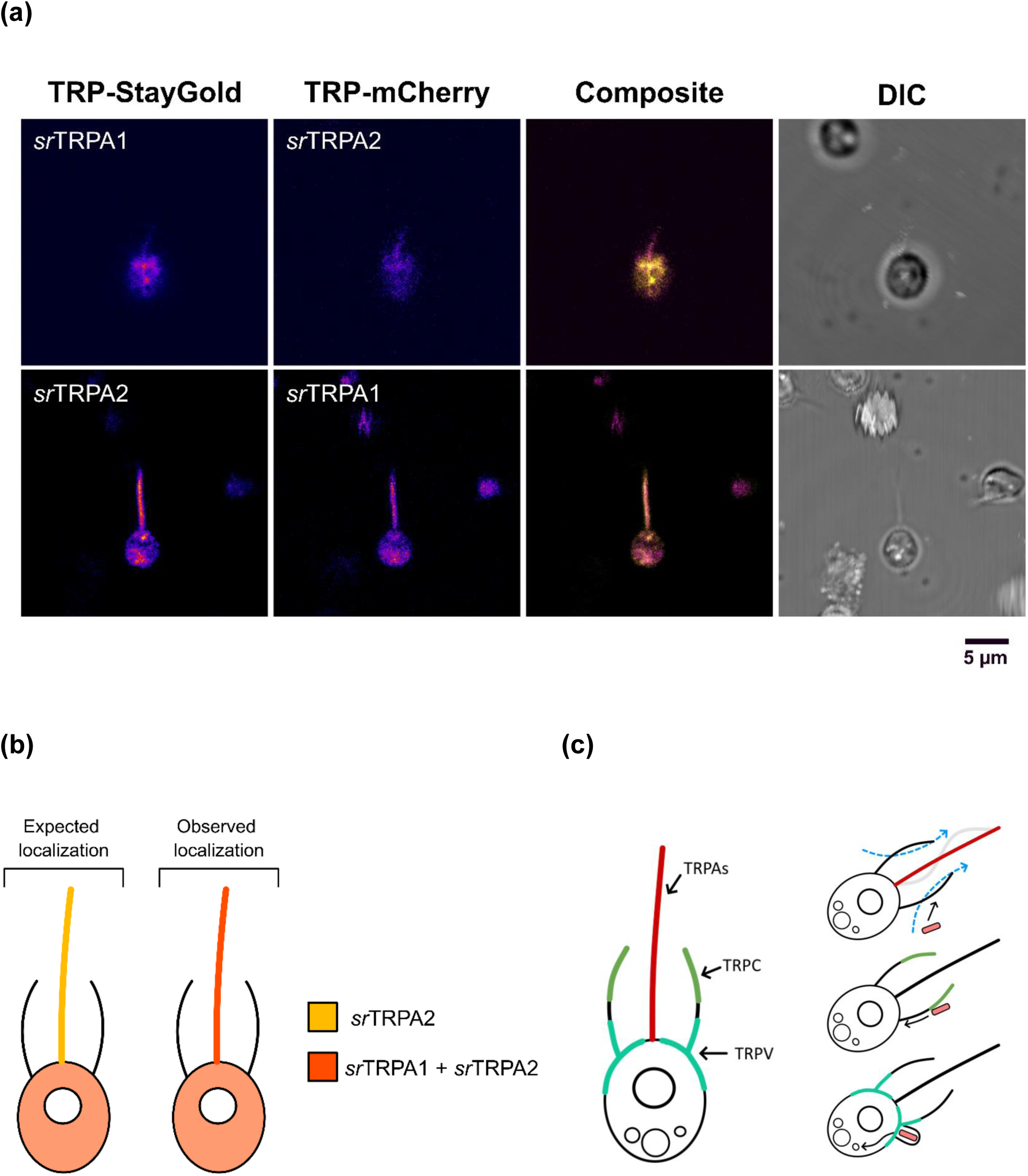
Co-overexpression of fluorescently tagged *sr*TRPA1 and *sr*TRPA2. **(a)** *sr*TRPA1 and *sr*TRPA2 were overexpressed in the same cells, coupled with either StayGold or mCherry, and imaged using confocal microscopy 1 day post-transfection. A differential interference contrast (DIC) image was used to observe the cell morphology. When co-overexpressed, both paralogs were observed in the flagellum. **(b)** Cartoon figures depicting the expected localization patterns for *sr*TRPA1 and *sr*TRPA2 based on single overexpression, and the observed patterns during co-overexpression. *sr*TRPA1 appears to require *sr*TRPA2 for flagellar localization. Similar colocalization was not observed when any of the TRPAs were co-overexpressed with *sr*TRPC or *sr*TRPV. **(c)** The three families of TRPA, TRPC and TRPV localize to distinct regions in the collar complex of *S. rosetta*. Each of these regions serve different roles in bacterial capture and feeding (see Figure 1a for details).

## Discussion

Our analysis of the TRP channel families in choanoflagellates and other non-animal holozoans is consistent with previous reports suggesting the presence of Group 1 and Group 2 families in the stem holozoan, while identifying a complex evolutionary history of the superfamily with the holozoan clade and pointing to strong functionalization in the choanoflagellate-animal ancestor (**Figure 3c**). There was then an expansion of Group 1 families leading to the choanoflagellate-animal ancestor, without large expansions within families, indicated by the lack of one-to-one orthologs between animals and choanoflagellates. After the divergence of choanoflagellates and animals, the TRP channel families of choanoflagellates appear to adopt two distinct patterns: TRPM and TRPW have undergone extensive radiations and appear to be plastic in choanoflagellate evolution; with many species-specific divergences, losses and lack of predictability between phylogenetic placement and the presence of domains such as SLOG and WD40 repeat domains. All other families remain restricted in numbers, and have more predictable domain organization, shorter branch lengths and correspondence between phylogenetic relationships of orthologs and of species.

The topology of the molecular phylogenies using the transmembrane and putative pore-forming regions suggests that the TRPW family do belong to the TRP channels, and appear to be monophyletic within choanoflagellates. However, it should be noted that there is some divergence within these regions and therefore the functionality as an ion channel will need to be experimentally confirmed. WD40 repeat domains are amongst the most common eukaryotic protein domains, typically forming 7-blade ꞵ-propeller structures that act as scaffolds for protein-protein interactions. The structural stability of these domains permits substantial sequence variation while maintaining folding integrity, facilitating evolutionary divergence of interaction partners (Hu et al., 2017; Xu & Min, 2011). In choanoflagellates, these domains could potentially recruit downstream signaling components to the site of channel activity. The presence of a TRPW in three of the four filastereans examined suggests that this family arose in at least the choanoflagellate-filasterean ancestor, and was subsequently lost in the stem animal group. Among choanoflagellates, multiple members of the family can be found in both Craspedida and Acanthoecida, suggesting early radiations within the clade. This family, along with TRPM, show higher and more variable numbers across the choanoflagellate lineages investigated, which could potentially result from a close connection with lifestyle, strengthening the ability to delineate variable inputs, in the absence of specialized cell types. These will be important targets for future studies in understanding choanoflagellate sensory biology.

In terms of subcellular localization, it is intriguing that all channels investigated (*sr*TRPA1, *sr*TRPA2, *sr*TRPC, & *sr*TRPV) localize to parts of the collar complex. The collar complex both generates the feeding current (via flagellar beating) and acts as the site of microbial capture. This mechanism results in interaction with solutes from a relatively large surrounding space (Sørensen et al., 2021), making it a likely site for detection of environmental signals. Furthermore, each of the different families appears to have a distinct domain of localization, with TRPAs localizing to the flagella, *sr*TRPC towards the tip of the microvillar collar, and *sr*TRPV at the collar base. This subcellular localization suggests distinct functions within the collar complex and points to a role in nuanced signaling in this part of the cell. These sub-complex domains each have specific roles in feeding, through generation of feeder current, bacterial capture, and phagocytosis (**Figure 5c**). The specific localization of different channels in each of these domains would be consistent with a role in regulating each of these processes.

TRP channels could transduce sensory input in the collar complex into electrical signals, which have previously been shown to elicit changes in flagellar beating and feeding behavior (Colgren & Burkhardt, 2025). In this scenario, the TRP channels are not necessarily the primary transducer, as animal TRP channels are often activated downstream of other sensory receptors such as GPCRs (Cox et al., 2024; Luo et al., 2025; Zhang et al., 2023) and could function in integrating signals that directly affect TRP channel gating with signals from other receptors. This creates a large combinatorial space around the core receptors to tune responses across a diverse range of stimuli. Focusing only on the single-cell state, there are conceptually a limited number of ways that single cells could delineate diverse environmental stimuli, including increased number of receptors, control of spatiotemporal deployment of the receptors, and increased interaction partners with modulatory potential. These are not mutually exclusive, and we see direct evidence for the first two in choanoflagellates, both with the radiations in TRPM and TRPW families as well as the discrete localization zones for TRPA, TRPC, and TRPV families.

Previous work has indicated the presence of several TRP channel families in the last common ancestor of animals and choanoflagellates (Cai, 2008; Cai & Clapham, 2012; Davila-Velderrain & van Giesen, 2025; Iordanov et al., 2019; Peng et al., 2015; Sigg et al., 2017), and our findings expand this picture, with TRPA, TRPC, TRPV, TRPVL, TRPM, TRPML, TRPP, and TRPW all predating the split between animals and choanoflagellates. There was likely a loss of the TRPW family in the stem animal lineage, as well as the emergence of TRPN channels likely derived from TRPC-like ancestors. These patterns suggest the evolution of animal sensory systems involved both pruning and innovation within an already complex ancestral toolkit (Colgren & Burkhardt, 2026).

Although the precise cellular organization of the first animals remains uncertain, it is likely that they were bacterivorous and may have possessed collar-type cells morphologically similar to those of modern choanoflagellates and sponges (Brunet & King, 2017; Laundon et al., 2019). In this case, there would be strong selective pressure towards the ability to resolve a heterogeneous microbial environment. Our observations that distinct TRP channel families are spatially segregated within the collar complex of *S. rosetta* suggest that these single cells can achieve functional diversification of sensory input through precise subcellular patterning of receptors. This further implies that mechanisms for spatial organization of sensory receptors predate the origin of animals and could have been co-opted for specialization of cell types during the transition to multicellularity. In choanoflagellates, diversification of sensory capacity appears to occur through a combination of receptor family expansion and subcellular specialization. With the advent of obligate multicellularity, these same principles could have been extended to the level of tissues and cell types, enabling the partitioning of sensory functions across distinct cellular populations.

In broader terms, the spatial segregation of TRP channels observed in modern choanoflagellates may represent an ancestral strategy for organizing sensory inputs that was later elaborated into the cell-type-specific expression patterns characteristic of animal sensory systems. The transition from subcellular to organismal patterning would thus not require entirely new molecular innovations, but rather a redeployment of pre-existing mechanisms for receptor localization and integration (Colgren & Burkhardt, 2026). Our findings are consistent with a model in which the evolution of animal sensory complexity was driven not only by the expansion of receptor repertoires, but also by changes in how these receptors are spatially and functionally organized. Understanding the molecular basis for establishing the different localization domains observed in modern choanoflagellates may provide essential insights into how specialization occurred in the first animal cell types.

## Acknowledgements

We thank all members of the Burkhardt group for their valuable insights and discussions, with particular thanks to Ruth Styfhals and Roberto Santoro for detailed feedback on the manuscript. We also thank Lena van Giesen and Fabian Rentzsch for their comments on the thesis that contributed to this work.

## Materials and Methods

### Identification and Annotation of *S. rosetta* TRP channels

52 protein sequences of well-annotated TRP channels from *Homo sapiens*, *Drosophila melanogaster* and *Caenorhabditis elegans* were downloaded from UniProtKB (**Supplemental file S1**). DeepTMHMM (Hallgren et al., 2022) was used to identify the transmembrane helices and the sequences were trimmed to contain the region delimited by the residues 50 residues directly upstream to 50 residues directly downstream of the TMD. This list was used as an entry in an PSI-BLAST search (Altschul et al., 1990) in the *S. rosetta* genome assembly GCA_000188695.1 (Fairclough et al., 2013; Yates et al., 2022). The search was done in three iterations with max numbers of sequences set to 500, 5000 and 10000, respectively. An E value of 10^-5^ was set as a threshold for hits to be included. All candidate TRP channels were subjected to a reciprocal BLASTP search and any candidates with non-TRP ion channels as the first hit were discarded.

DeepTMHMM and InterProScan (Jones et al., 2014) were used to examine all candidate sequences, and AlphaFold 3 (Abramson et al., 2024) was used to predict tetrameric structures for most candidates, although some were analyzed as monomers due to limitations in AlphaFold. When domain structure was predicted with high confidence levels, the same domain was retrieved from a protein of known structure and superimposed on the predicted structure using TMalign (Zhang & Skolnick, 2005) to look for deviations. MUSCLE (Edgar, 2004) was also used to create multiple sequence alignments (MSAs) with metazoan homologs (if any) to look for conserved features. Together with a phylogenetic tree, these approaches culminated in manual annotation of the sequences.

### Constructing Phylogenetic Trees

The TM predictions were used to trim all sequences to include the region beginning 50 aa upstream of the first TM helix and ending 50 aa after the last TM helix. If more than six TM helices were present, only the last six were considered (in line with InterProScan predictions for the *S. rosetta* candidates and metazoan homologs). The sequences were aligned using MAFFT (Katoh & Standley, 2013) (standard parameters) and a maximum-likelihood phylogenetic tree was constructed using IQ-tree 2 (Minh et al., 2020). The integrated ModelFinder algorithm was used to select the substitution model. Both UFBoot (Minh et al., 2013) and SH-aLRT (Guindon et al., 2010) Bootstrapping was done with 1000 replicates. Trees were visualized in iTOL (Letunic & Bork, 2021).

### Searching for and Annotating Orthologs

The TMD and 50 flanking residues of all candidate *S. rosetta* TRP channels, as well as the metazoan homologs used in the initial BLAST search to identify them, were used to create a hidden Markov model (HMM) using HMMER (Potter et al., 2018) to identify orthologs in other species. The HMM was tested on a well-annotated genome-derived proteome of mouse (Ensembl: GRCm38) and the resulting E values were used to set 10^-20^ as the threshold for hits to be included. Although some of the included hits were VGICs, with E values towards the upper limit, subsequent filtering by DeepTMHMM-predicted TM topology would remove them due to differing TM topology. The HMM was applied to 19 choanoflagellate transcriptome-derived and one genome-derived proteomes (Brunet, 2019; King et al., 2008; Richter, 2018), as well as 10 genome-derived proteomes for non-choanozoan holozoans (sequences downloaded from MultiCellGenome Lab: https://multicellgenome.com/genomes).

The Pfam-A set of HMMs (Mistry et al., 2021) was applied to search for the following domains: WD40 (PF00400.38), Ion_trans (PF00520.37), LSDAT_euk (PF18139.7) (SLOG), NUDIX (PF00293.34), Ank (PF00023.36) and Isochorismatase (PF00857.26) for all sequences. To search for the extracellular domain of TRPPs, a custom HMM was built based on the S1-S2 extracellular region of animal reference sequences, using HMMer as described above.

### Constructing Sequence Similarity Network (SSN)

All full-length sequences (both those of interest, and others used for reference) were used as query in an all-against-all BLAST search (Camacho et al., 2009) and edges with E values less than 10^-20^ were imported into Cytoscape (Shannon et al., 2003) for visualization. Some nodes were moved to more easily see relevant edges, and any singular nodes were added manually to the figure. To annotate the TRPVL cluster, a TRPVL from *Nematostella vectensis* (jgi|Nemve1|21409) was used as query in a BLAST search among all the choanoflagellate TRP channels, giving clear clustering with the members of the TRPVL cluster.

### Examining residue-specific conservation

Residue-specific conservation was examined within the TRP channel families using two methods in parallel. In the first method an *S. rosetta* member with a generated AlphaFold structure was chosen and the full-length sequences were aligned by MAFFT, and positions in the MSA where the reference had gaps were discarded. The rate4site (Pupko et al., 2002) was used to calculate conservation, and these values were subsequently projected onto the structure, followed by manual inspection.

To account for positions of the MSA lost in the above approach, all sequences were aligned and the Shannon entropy was calculated for each position in the MSA. These values were inverted and normalized to a 0 to 1 scale corresponding to low to high conservation, and visualized together with occupancy at each position of the MSA, and whether the reference sequence was present at that position.

### Culturing S. rosetta

*Salpingoeca rosetta* was cocultured with the bacterium *Echinicola pacifica* in 6 mL high nutrient media (HNM) at 22°C in a sterile environment. The cells were passaged every 2-3 days in 1:60 or 1:120 dilutions, depending on apparent cell density. The cultures were vigorously shaken before passaging. In case of excessive *E. pacifica* growth, the cultures were washed by one or more rounds of centrifugation at 2000 rcf for 5 minutes and resuspension of the pellet.

### Plasmid Design

The plasmids were designed to encode a fusion protein consisting of the TRP channel of interest and a fluorophore attached to the C-terminus by a flexible linker region. The plasmid NK802 was used as the backbone and included a puromycin-resistance gene (pac) driven by the promoter pEF1 (elongation factor 1) derived from *S. rosetta*. The TRP channel insert was amplified from cDNA, while the fluorophore StayGold was amplified from a synthetic construct that was codon optimized for *S. rosetta* (codon usage based on (Southworth et al., 2018)). When later creating the TRP-mCherry constructs, the same approach as described below was used, but NK802 already contains mCherry, and only had to be linearized using PCR (see **Table S1** for complete list of primers).

### cDNA Synthesis and Amplification

cDNA was synthesized from total RNA previously derived from *S. rosetta* rosette colonies. 2.5 μL of total RNA was mixed with oligo(dT)_20_ and dNTPs to final concentrations of 3.8 μM and 0.77 μM, respectively, in a total volume of 13 μL. The mixture was heated to 65°C for 5 minutes, before immediately placing it on ice. This mixture was incorporated into a 20 μL reaction mixture that was made to also contain 1x First-Strand Buffer, 5 mM DTT, 40 U RNaseOUTTM Recombinant RNase Inhibitor, and 200 U SuperScriptTM III Reverse Transcriptase. The reaction mixtures were incubated at 50°C for 60 minutes, followed by 70°C for 15 minutes to deactivate the reaction.

PCR was used to amplify the coding regions of cDNA and to add additional homology arms at each end to facilitate plasmid integration. The primers used for cDNA amplification were given in **Table S1** and each primer had a 30 nt 5’ overhang complementary to the plasmid. Additionally, the reverse primers contained a region to serve as a flexible linker region between the protein of interest and fluorophore. For each target gene, 50 μL PCR mixtures were made containing 0.50 μM of both gene-specific primers, 1x Q5^®^ Reaction Buffer, 0.20 mM dNTPs, 0.4 μL of the first-strand reverse transcription product, and 0.5 U Q5^®^ High-Fidelity DNA Polymerase. A 35-cycle PCR program was run with denaturation at 98°C for 10 s, annealing at 65°C for 30 s and extension at 72°C for 150 s. The program was initiated by an additional denaturation step for 30 s and terminated by a 120 s final elongation. The PCR products were separated by agarose gel electrophoresis and subsequently the band of interest was excised and purified using the Monarch^®^ DNA Gel Extraction Kit according to its accompanying guidelines.

### Plasmid synthesis

PCR was used to amplify a section of the plasmid NK802 into a linear fragment to serve as a backbone for subsequent cloning, and to amplify the StayGold coding sequence from a synthetic construct. The primers GAGCTCCCCCCAGCATTATCACG and GGCTGGTTGTTTTGTGGTTGTGG, and ATGGCCTCCACCCCCTTCAAG and GAGGTGGGCCTCGAGGGTC were used for the plasmid backbone and StayGold, respectively. For each, two 50 μL reaction mixtures were made containing 0.5 μM of each of the appropriate forward and reverse primer, 1x Phusion™ HF Buffer, 200 μM dNTPs, and 1 U Phusion™ High-Fidelity DNA Polymerase. 1 ng of template was used per reaction, and the. The same PCR program as above was used for 40 cycles, although annealing temperature was 68°C and elongation times were 210 s and 30 s for the linearized plasmid and StayGold, respectively. Residual plasmid template was digested by DpnI (1 U) at 37°C for 1 hour, and both products were subsequently purified by gel extraction as before.

The linearized plasmid NK802, StayGold insert, gene-specific insert and a linker region to promote proper assembly were mixed at molar ratios of 1:2:2:3 and diluted to final volumes of 5 μL or 10 μL depending on the amount of linearized plasmid available. The linker region (CCAGTCCGAGACCCTCGAGGCCCACCTCTAAGAGCTCCCCCCAGC-ATTATCACGTCTACT) was made by mixing equimolar amounts of complementary ssDNA sequences and heating to 95°C for 5 minutes before allowing it to slowly cool to RT. The reaction mixtures were made by adding 2x NEB HiFi DNA Assembly Master Mix to an equal volume of the fragment mixtures. The mixtures were incubated at 50°C for 1 hour to assemble the plasmids

NEB 5α competent E. coli cells were thawed on ice, and 2 μL of assembled plasmid products were added to the bacteria. The cells were left on ice for 30 minutes before heat shocking at 42°C for 30 s and putting them back on ice for 5 minutes. This was followed by incubation in 950 μL of S.O.C. medium at 37°C under vigorous shaking for 60 minutes. After incubation, 100 μL of the bacterial culture was added to LB agar plates with 80 μg mL^-1^ ampicillin, before incubating the plates at 37°C for 20 hours.

Colonies were then picked at random and dipped into 10 µL of TE buffer with 0.1% (v/v) Triton X-100 before heating the mixtures to 95°C for 10 minutes to lyse the bacteria. 2 μL of this mixture was used as template for PCR in 50 μL-reactions made to contain 1x Green GoTaq^®^ Flexi Buffer, 2.0 mM MgCl_2_, 0.2 mM dNTPs, 0.50 μM each of the primers CGTGATAATGCTGGGGGGAGCTC and CCACAACCACAAAACAACCAGCC, and 1.25 U of GoTaq^®^ G2 Flexi DNA Polymerase. A 35-cycle program with denaturation at 95°C for 30 s (initial cycle 150 s), annealing at 56°C for 30 s, and elongation at 72°C for 240 s (final elongation 840 s). Colonies with appropriate bands were selected for inoculation of LB medium containing 80 µg mL^-1^ ampicillin and incubated overnight at 37°C during vigorous shaking, and plasmids were purified using the Qiagen mini prep kit according to the accompanying instructions. Part of the cultures were cryopreserved in 25% (v/v) glycerol and stored at -70°C.

### Sanger sequencing of plasmids was done externally by Genewiz

Once proper assembly had been validated by sequencing, the preserved cultures were used to inoculate 100 mL ampicillin-containing LB medium, incubated overnight and harvested using Qiagen Midi Prep kit according to the given instructions and concentrated to 5.0 µg µL^-1^ using DNA precipitation.

### Transfection of *S. rosetta*

The transfection methodology for S. rosetta was based on that of (Booth et al., 2018). *S. rosetta* cultures were seeded at 8000 cells mL^-1^ in 50 mL HNM about 48 hours prior to transfection. The cultures were inspected on the day of transfection to ensure that the cells were in the slow swimmer and chain states. The cells were then first washed by vigorously shaking the culture, centrifuging at 2000 rcf at RT for 5 minutes, removing the supernatant, and resuspending in 50 mL artificial seawater (ASW). This washing step was repeated, before having an additional centrifugation at 2200 rcf for 5 minutes and removing as much of the supernatant as possible. The pellet was then resuspended in 250 μL of ASW and the cell density was calculated by first taking out 2 μL of this suspension, diluting by a factor of 100 in ASW, and adding formaldehyde to a final concentration of 0.37% (v/v) to fix the cells. Cell counting was done by the Luna-FLTM Dual Fluorescence Cell Counter. This was followed by diluting to 5·10^7^ cells mL^-1^ and splitting this into 100 μL aliquots. One of these was centrifugated at 800 rcf at RT for 5 minutes. The pellet was resuspended in 100 μL freshly prepared priming buffer containing 3 μM papain and was incubated at RT for 35 minutes to degrade the glycocalyx of the cells. BSA was added to the cells at a final concentration of 4.5 mg mL^-1^ to quench papain digestion, followed by centrifugation at 1250 rcf at RT for 5 minutes. The cell pellet was resuspended in 25 μL of ice-cold SF buffer and kept on ice until nucleofection. The nucleofection mixtures were made by mixing 1 μL of each of the plasmids at 5 mg mL^-1^, 2 μL of pUC19 at 20 mg mL^-1^ as carrier plasmid, 16 μL of ice-cold SF buffer and 2 μL of the cell suspension. The mixtures were transferred to nucleofection cuvettes and inserted into a 4D-Nucleofector. Pulsing was done using the CM156 pulse and was followed by a swift addition of 100 μL of ice-cold recovery buffer. After 10 minutes at RT, the cells were transferred to 2 mL low nutrient media (LNM) and placed at 22°C. After 1 hour, 0.1 mg of *E. pacifica* was added to feed the choanoflagellates.

### Confocal Microscopy

The first imaging of the transfected cells was done 20-28 hours after pulsing. For imaging, 1 mL of the cultures were washed by centrifugation at 2000 rcf at 4°C for 10 minutes before resuspending the pellet in 1 mL of ASW and repeating the centrifugation. The cells were then resuspended in 150 μL of ASW and plated on poly-L-lysine-coated glass-bottom dishes and left to settle for 30 minutes before imaging.

Imaging was performed on an Olympus FV3000RS confocal microscope using FLUOVIEW acquisition software (Olympus). Images were acquired using a 60x objective (Olympus; UPlanSApo 60x/1.30 Sil) or 100x objective (Olympus; UPlanSApo 100x/1.40 Oil). Cells were imaged using preloaded settings for AZAMI green and mCherry. The laser intensities during imaging were adjusted to exclude autofluorescence signal in non-transfected cells. The cells were imaged less than 2.5 hours after plating due to a decrease in viability during imaging.

### Puromycin Selection and Clonal Isolation

Puromycin was added to the part of the cultures not used for imaging on the first day at a final concentration of 40 μg mL^-1^ to select for cells that maintained the plasmid. The imaging was repeated 5 days after transfection as described above. For cells transfected with *sr*TRPA1-StayGold and *sr*TRPA2-StayGold plasmids, puromycin resistance persisted and monoclonal cultures were obtained by dilution into 3 cells mL^-1^ and passaged into 0.1 mL subcultures. Cultures for both were cryo-preserved (Chandra & Rutaganira, 2025), but the proportion of fluorescent cells remained low despite the isolation into monoclonal cultures.

## References

1. Abe, K., & Puertollano, R. (2011). Role of TRP channels in the regulation of the endosomal pathway. Physiology (Bethesda), 26(1), 14–22. 10.1152/physiol.00048.2010

2. Abramson, J., Adler, J., Dunger, J., Evans, R., Green, T., Pritzel, A., Ronneberger, O., Willmore, L., Ballard, A. J., Bambrick, J., Bodenstein, S. W., Evans, D. A., Hung, C.-C., O’Neill, M., Reiman, D., Tunyasuvunakool, K., Wu, Z., Žemgulytė, A., Arvaniti, E., Jumper, J. M. (2024). Accurate structure prediction of biomolecular interactions with AlphaFold 3. Nature, 630(8016), 493–500. 10.1038/s41586-024-07487-w

3. Ahmed, T., Nisler, C. R., Fluck, E. C., 3rd, Walujkar, S., Sotomayor, M., & Moiseenkova-Bell, V. Y. (2022). Structure of the ancient TRPY1 channel from Saccharomyces cerevisiae reveals mechanisms of modulation by lipids and calcium. Structure, 30(1), 139–155.e135. 10.1016/j.str.2021.08.003

4. Altschul, S. F., Gish, W., Miller, W., Myers, E. W., & Lipman, D. J. (1990). Basic Local Alignment Search Tool. Journal of Molecular Biology, 215(3), 403–410. 10.1006/jmbi.1990.9999

5. Arendt, D. (2021). Elementary nervous systems. Philos Trans R Soc Lond B Biol Sci, 376(1821), 20200347. 10.1098/rstb.2020.0347

6. Bellemer, A. (2015). Thermotaxis, circadian rhythms, and TRP channels in Drosophila. Temperature (Austin), 2(2), 227–243. 10.1080/23328940.2015.1004972

7. Booth, D. S., Szmidt-Middleton, H., & King, N. (2018). Transfection of choanoflagellates illuminates their cell biology and the ancestry of animal septins. Mol Biol Cell, 29(25), 3026–3038. 10.1091/mbc.E18-08-0514

8. Brunet, T. (2019). Choanoeca flexa transcriptome and predicted nonredundant proteome. 10.6084/m9.figshare.5686984.v2

9. Brunet, T., Albert, M., Roman, W., Coyle, M. C., Spitzer, D. C., & King, N. (2021). A flagellate-to-amoeboid switch in the closest living relatives of animals. Elife, 10, e61037. 10.7554/eLife.61037

10. Brunet, T., & King, N. (2017). The Origin of Animal Multicellularity and Cell Differentiation. Dev Cell, 43(2), 124–140. 10.1016/j.devcel.2017.09.016

11. Brunet, T., Larson, B. T., Linden, T. A., Vermeij, M. J. A., McDonald, K., & King, N. (2019). Light-regulated collective contractility in a multicellular choanoflagellate. Science, 366(6463), 326–334. 10.1126/science.aay2346

12. Cai, X. (2008). Unicellular Ca2+ signaling ’toolkit’ at the origin of metazoa. Mol Biol Evol, 25(7), 1357–1361. 10.1093/molbev/msn077

13. Cai, X., & Clapham, D. E. (2012). Ancestral Ca2+ signaling machinery in early animal and fungal evolution. Mol Biol Evol, 29(1), 91–100. 10.1093/molbev/msr149

14. Camacho, C., Coulouris, G., Avagyan, V., Ma, N., Papadopoulos, J., Bealer, K., & Madden, T. L. (2009). BLAST+: architecture and applications. BMC Bioinformatics, 10(1), 421. 10.1186/1471-2105-10-421

15. Carr, M., Leadbeater, B. S. C., Hassan, R., Nelson, M., & Baldauf, S. L. (2008). Molecular phylogeny of choanoflagellates, the sister group to Metazoa. Proceedings of the National Academy of Sciences of the United States of America, 105(43), 16641–16646. 10.1073/pnas.0801667105

16. Carr, M., Richter, D. J., Fozouni, P., Smith, T. J., Jeuck, A., Leadbeater, B. S. C., & Nitsche, F. (2017). A six-gene phylogeny provides new insights into choanoflagellate evolution. Molecular Phylogenetics and Evolution, 107, 166–178. 10.1016/j.ympev.2016.10.011

17. Chandra, S., & Rutaganira, F. U. (2025). Glycerol improves the viability of a cryopreserved choanoflagellate. Cryobiology, 118, 105183. 10.1016/j.cryobiol.2024.105183

18. Colgren, J., & Burkhardt, P. (2025). Electrical signaling and coordinated behavior in the closest relative of animals. Sci Adv, 11(2), eadr7434. 10.1126/sciadv.adr7434

19. Colgren, J. J., & Burkhardt, P. (2026). The evolutionary origins of synaptic proteins and their changing roles in different organisms across evolution. Nat Rev Neurosci, 27(1), 7–22. 10.1038/s41583-025-00983-6

20. Cox, C. D., Poole, K., & Martinac, B. (2024). Re-evaluating TRP channel mechanosensitivity. Trends Biochem Sci, 49(8), 693–702. 10.1016/j.tibs.2024.05.004

21. Davila-Velderrain, J., & van Giesen, L. (2025). Voltage-gated ion channel diversity underlies neuronal excitability and nervous system evolution. Nat Commun, 16(1), 11534. 10.1038/s41467-025-66641-8

22. Dayel, M. J., Alegado, R. A., Fairclough, S. R., Levin, T. C., Nichols, S. A., McDonald, K., & King, N. (2011). Cell differentiation and morphogenesis in the colony-forming choanoflagellate Salpingoeca rosetta. Dev Biol, 357(1), 73–82. 10.1016/j.ydbio.2011.06.003

23. Dayel, M. J., & King, N. (2014). Prey capture and phagocytosis in the choanoflagellate Salpingoeca rosetta. PLoS One, 9(5), e95577. 10.1371/journal.pone.0095577

24. Edgar, R. C. (2004). MUSCLE: multiple sequence alignment with high accuracy and high throughput. Nucleic Acids Res, 32(5), 1792–1797. 10.1093/nar/gkh340

25. Esposito, S. L. J. L. M. E. K. J. (2021). Evolution of the Sensory/Neural Cell Types. In Origin and Evolution of Metazoan Cell Types (1 ed.). CRC Press.

26. Fairclough, S. R., Chen, Z., Kramer, E., Zeng, Q., Young, S., Robertson, H. M., Begovic, E., Richter, D. J., Russ, C., Westbrook, M. J., Manning, G., Lang, B. F., Haas, B., Nusbaum, C., & King, N. (2013). Premetazoan genome evolution and the regulation of cell differentiation in the choanoflagellate Salpingoeca rosetta. Genome Biol, 14(2), R15. 10.1186/gb-2013-14-2-r15

27. Garcia De Las Bayonas, A., & King, N. (2025). G-protein-coupled receptor diversity and evolution in the closest living relatives of metazoa. Elife, 14, RP107467. 10.7554/eLife.107467

28. Glockling, S. L., Marshall, W. L., & Gleason, F. H. (2013). Phylogenetic interpretations and ecological potentials of the Mesomycetozoea (Ichthyosporea). Fungal Ecology, 6(4), 237–247. 10.1016/j.funeco.2013.03.005

29. Grau-Bové, X., Torruella, G., Donachie, S., Suga, H., Leonard, G., Richards, T. A., & Ruiz-Trillo, I. (2017). Dynamics of genomic innovation in the unicellular ancestry of animals. Elife, 6, e26036. 10.7554/eLife.26036

30. Guindon, S., Dufayard, J. F., Lefort, V., Anisimova, M., Hordijk, W., & Gascuel, O. (2010). New algorithms and methods to estimate maximum-likelihood phylogenies: assessing the performance of PhyML 3.0. Syst Biol, 59(3), 307–321. 10.1093/sysbio/syq010

31. Hallgren, J., Tsirigos, K. D., Pedersen, M. D., Almagro Armenteros, J. J., Marcatili, P., Nielsen, H., Krogh, A., & Winther, O. (2022). DeepTMHMM predicts alpha and beta transmembrane proteins using deep neural networks. bioRxiv, 2022.2004.2008.487609. 10.1101/2022.04.08.487609

32. Himmel, N. J., & Cox, D. N. (2020). Transient receptor potential channels: current perspectives on evolution, structure, function and nomenclature. Proc Biol Sci, 287(1933), 20201309. 10.1098/rspb.2020.1309

33. Himmel, N. J., Gray, T. R., & Cox, D. N. (2020). Phylogenetics Identifies Two Eumetazoan TRPM Clades and an Eighth TRP Family, TRP Soromelastatin (TRPS). Molecular Biology and Evolution, 37(7), 2034–2044. 10.1093/molbev/msaa065

34. Hsiao, J., Deng, L. C., Chalasani, S., & Edsinger, E. (2021). Numerous expansions in TRP ion channel diversity highlight widespread evolution of molecular sensors in animal diversification. bioRxiv, 2021.2011.2014.466824. 10.1101/2021.11.14.466824

35. Hu, X.-J., Li, T., Wang, Y., Xiong, Y., Wu, X.-H., Zhang, D.-L., Ye, Z.-Q., & Wu, Y.-D. (2017). Prokaryotic and Highly-Repetitive WD40 Proteins: A Systematic Study. Scientific Reports, 7(1), 10585. 10.1038/s41598-017-11115-1

36. Iordanov, I., Tóth, B., Szollosi, A., & Csanády, L. (2019). Enzyme activity and selectivity filter stability of ancient TRPM2 channels were simultaneously lost in early vertebrates. Elife, 8, e44556. 10.7554/eLife.44556

37. Ireland Ella, V., Woznica, A., & King, N. (2020). Synergistic Cues from Diverse Bacteria Enhance Multicellular Development in a Choanoflagellate. Applied and Environmental Microbiology, 86(11), e02920–02919. 10.1128/AEM.02920-19

38. Jaślan, D., Böck, J., Krogsaeter, E., & Grimm, C. (2020). Evolutionary Aspects of TRPMLs and TPCs. Int J Mol Sci, 21(11). 10.3390/ijms21114181

39. Jones, P., Binns, D., Chang, H.-Y., Fraser, M., Li, W., McAnulla, C., McWilliam, H., Maslen, J., Mitchell, A., Nuka, G., Pesseat, S., Quinn, A. F., Sangrador-Vegas, A., Scheremetjew, M., Yong, S.-Y., Lopez, R., & Hunter, S. (2014). InterProScan 5: genome-scale protein function classification. Bioinformatics, 30(9), 1236–1240. 10.1093/bioinformatics/btu031

40. Julius, D. (2013). TRP channels and pain. Annu Rev Cell Dev Biol, 29, 355–384. 10.1146/annurev-cellbio-101011-155833

41. Karapidaki, I., Handberg-Thorsager, M., Momose, T., Yasuo, H., Genikhovich, G., Assaf, S., Deleau, C., Pang, Y., Pavlich, C., Lohmann, B., Rusciano, M. L., Stranges, M., Mathieu, J., Zilliox, M., Ustyantsev, K., Salmon, B., Laplace-Builhé, B., Koenig, M., Colgren, J. J., . . . Averof, M. (2026). Targeting the cell membrane in established and emerging model organisms. Development, 153(8). 10.1242/dev.205415

42. Kashio, M., & Tominaga, M. (2022). TRP channels in thermosensation. Curr Opin Neurobiol, 75, 102591. 10.1016/j.conb.2022.102591

43. Katoh, K., & Standley, D. M. (2013). MAFFT Multiple Sequence Alignment Software Version 7: Improvements in Performance and Usability. Molecular Biology and Evolution, 30(4), 772–780. 10.1093/molbev/mst010

44. King, N., Westbrook, M. J., Young, S. L., Kuo, A., Abedin, M., Chapman, J., Fairclough, S., Hellsten, U., Isogai, Y., Letunic, I., Marr, M., Pincus, D., Putnam, N., Rokas, A., Wright, K. J., Zuzow, R., Dirks, W., Good, M., Goodstein, D., Rokhsar, D. (2008). The genome of the choanoflagellate Monosiga brevicollis and the origin of metazoans. Nature, 451(7180), 783–788. 10.1038/nature06617

45. Kirkegaard, J. B., Bouillant, A., Marron, A. O., Leptos, K. C., & Goldstein, R. E. (2016). Aerotaxis in the closest relatives of animals. Elife, 5, e18109. 10.7554/eLife.18109

46. Laundon, D., Larson, B. T., McDonald, K., King, N., & Burkhardt, P. (2019). The architecture of cell differentiation in choanoflagellates and sponge choanocytes. PLoS Biol, 17(4), e3000226. 10.1371/journal.pbio.3000226

47. Laursen, W. J., Anderson, E. O., Hoffstaetter, L. J., Bagriantsev, S. N., & Gracheva, E. O. (2015). Species-specific temperature sensitivity of TRPA1. Temperature (Austin), 2(2), 214–226. 10.1080/23328940.2014.1000702

48. Leadbeater, B. S. C. (2015). The Choanoflagellates: Evolution, Biology and Ecology. Cambridge University Press. 10.1017/CBO9781139051125

49. Letunic, I., & Bork, P. (2021). Interactive Tree Of Life (iTOL) v5: an online tool for phylogenetic tree display and annotation. Nucleic Acids Research, 49(W1), W293–W296. 10.1093/nar/gkab301

50. Liao, M., Cao, E., Julius, D., & Cheng, Y. (2013). Structure of the TRPV1 ion channel determined by electron cryo-microscopy. Nature, 504(7478), 107–112. 10.1038/nature12822

51. Lindström, J. B., Pierce, N. T., & Latz, M. I. (2017). Role of TRP Channels in Dinoflagellate Mechanotransduction. Biol Bull, 233(2), 151–167. 10.1086/695421

52. Liu, H., Steenwyk, J. L., Zhou, X., Schultz, D. T., Kocot, K. M., Shen, X. X., Rokas, A., & Li, Y. (2024). A taxon-rich and genome-scale phylogeny of Opisthokonta. PLoS Biol, 22(9), e3002794. 10.1371/journal.pbio.3002794

53. Lu, X., Yao, Z., Wang, Y., Yin, C., Li, J., Chai, L., Dong, W., Yuan, L., Lai, R., & Yang, S. (2022). The acquisition of cold sensitivity during TRPM8 ion channel evolution. Proc Natl Acad Sci U S A, 119(21), e2201349119. 10.1073/pnas.2201349119

54. Luo, Y., Sun, L., & Peng, Y. (2025). The structural basis of the G protein-coupled receptor and ion channel axis. Curr Res Struct Biol, 9, 100165. 10.1016/j.crstbi.2025.100165

55. Minh, B. Q., Nguyen, M. A. T., & von Haeseler, A. (2013). Ultrafast Approximation for Phylogenetic Bootstrap. Molecular Biology and Evolution, 30(5), 1188–1195. 10.1093/molbev/mst024

56. Minh, B. Q., Schmidt, H. A., Chernomor, O., Schrempf, D., Woodhams, M. D., von Haeseler, A., & Lanfear, R. (2020). IQ-TREE 2: New Models and Efficient Methods for Phylogenetic Inference in the Genomic Era. Molecular Biology and Evolution, 37(5), 1530–1534. 10.1093/molbev/msaa015

57. Miño, G. L., Koehl, M. A. R., King, N., & Stocker, R. (2017). Finding patches in a heterogeneous aquatic environment: pH-taxis by the dispersal stage of choanoflagellates. Limnology and Oceanography Letters, 2(2), 37–46. 10.1002/lol2.10035

58. Mistry, J., Chuguransky, S., Williams, L., Qureshi, M., Salazar, Gustavo A., Sonnhammer, E. L. L., Tosatto, S. C. E., Paladin, L., Raj, S., Richardson, L. J., Finn, R. D., & Bateman, A. (2021). Pfam: The protein families database in 2021. Nucleic Acids Research, 49(D1), D412–D419. 10.1093/nar/gkaa913

59. Nitsche, F., Carr, M., Arndt, H., & Leadbeater, B. S. (2011). Higher level taxonomy and molecular phylogenetics of the Choanoflagellatea. J Eukaryot Microbiol, 58(5), 452–462. 10.1111/j.1550-7408.2011.00572.x

60. Peng, G., Shi, X., & Kadowaki, T. (2015). Evolution of TRP channels inferred by their classification in diverse animal species. Mol Phylogenet Evol, 84, 145–157. 10.1016/j.ympev.2014.06.016

61. Potter, S. C., Luciani, A., Eddy, S. R., Park, Y., Lopez, R., & Finn, R. D. (2018). HMMER web server: 2018 update. Nucleic Acids Research, 46(W1), W200–W204. 10.1093/nar/gky448

62. Pupko, T., Bell, R. E., Mayrose, I., Glaser, F., & Ben-Tal, N. (2002). Rate4Site: an algorithmic tool for the identification of functional regions in proteins by surface mapping of evolutionary determinants within their homologues. Bioinformatics, 18(suppl_1), S71–S77. 10.1093/bioinformatics/18.suppl_1.S71

63. Richter, D., & Nitsche, F. (2017). Choanoflagellatea. In (pp. 1479–1496). 10.1007/978-3-319-28149-0_5

64. Richter, D. F., Parinaz; Eisen, Michael; King, Nicole (2018). Data from: Gene family innovation, conservation and loss on the animal stem lineage. 10.6084/m9.figshare.5686984.v2

65. Ros-Rocher, N., & Brunet, T. (2023). What is it like to be a choanoflagellate? Sensation, processing and behavior in the closest unicellular relatives of animals. Anim Cogn, 26(6), 1767–1782. 10.1007/s10071-023-01776-z

66. Schindl, R., & Romanin, C. (2007). Assembly domains in TRP channels. Biochem Soc Trans, 35(Pt 1), 84–85. 10.1042/bst0350084

67. Schlosser, G. (2025). Evolutionary History of Sensory Cell Types. In: Oxford University Press.

68. Schüler, A., Schmitz, G., Reft, A., Özbek, S., Thurm, U., & Bornberg-Bauer, E. (2015). The Rise and Fall of TRP-N, an Ancient Family of Mechanogated Ion Channels, in Metazoa. Genome Biol Evol, 7(6), 1713–1727. 10.1093/gbe/evv091

69. Sebé-Pedrós, A., Irimia, M., del Campo, J., Parra-Acero, H., Russ, C., Nusbaum, C., Blencowe, B. J., & Ruiz-Trillo, I. (2013). Regulated aggregative multicellularity in a close unicellular relative of metazoa. Elife, 2, e01287. 10.7554/eLife.01287

70. Shabardina, V., Dharamshi, J. E., Ara, P. S., Anto, M., Bascon, F. J., Suga, H., Marshall, W., Scazzocchio, C., Casacuberta, E., & Ruiz-Trillo, I. (2024). Ichthyosporea: a window into the origin of animals. Communications Biology, 7(1). https://doi.org/ARTN 915 10.1038/s42003-024-06608-5

71. Shannon, P., Markiel, A., Ozier, O., Baliga, N. S., Wang, J. T., Ramage, D., Amin, N., Schwikowski, B., & Ideker, T. (2003). Cytoscape: a software environment for integrated models of biomolecular interaction networks. Genome Res, 13(11), 2498–2504. 10.1101/gr.1239303

72. Sigg, M. A., Menchen, T., Lee, C., Johnson, J., Jungnickel, M. K., Choksi, S. P., Garcia, G., 3rd, Busengdal, H., Dougherty, G. W., Pennekamp, P., Werner, C., Rentzsch, F., Florman, H. M., Krogan, N., Wallingford, J. B., Omran, H., & Reiter, J. F. (2017). Evolutionary Proteomics Uncovers Ancient Associations of Cilia with Signaling Pathways. Dev Cell, 43(6), 744–762.e711. 10.1016/j.devcel.2017.11.014

73. Southworth, J., Armitage, P., Fallon, B., Dawson, H., Bryk, J., & Carr, M. (2018). Patterns of Ancestral Animal Codon Usage Bias Revealed through Holozoan Protists. Molecular Biology and Evolution, 35(10), 2499–2511. 10.1093/molbev/msy157

74. Startek, J. B., Boonen, B., Talavera, K., & Meseguer, V. (2019). TRP Channels as Sensors of Chemically-Induced Changes in Cell Membrane Mechanical Properties. Int J Mol Sci, 20(2). 10.3390/ijms20020371

75. Stocker, R. (2012). Marine Microbes See a Sea of Gradients. Science, 338(6107), 628–633. 10.1126/science.1208929

76. Sørensen, S., Asadzadeh, S. S., & Walther, J. H. (2021). Hydrodynamics of Prey Capture and Transportation in Choanoflagellates. Fluids, 6(3), 94.

77. Talyzina, I. A., Nadezhdin, K. D., & Sobolevsky, A. I. (2024). Forty sites of TRP channel regulation. Curr Opin Chem Biol, 84, 102550. 10.1016/j.cbpa.2024.102550

78. Valencia-Montoya, W. A., Pierce, N. E., & Bellono, N. W. (2024). Evolution of Sensory Receptors. Annu Rev Cell Dev Biol, 40(1), 353–379. 10.1146/annurev-cellbio-120123-112853

79. Wan, K. Y., & Jékely, G. (2021). Origins of eukaryotic excitability. Philos Trans R Soc Lond B Biol Sci, 376(1820), 20190758. 10.1098/rstb.2019.0758

80. Won, J., Kim, J., Kim, J., Ko, J., Park, C. H., Jeong, B., Lee, S. E., Jeong, H., Kim, S. H., Park, H., So, I., & Lee, H. H. (2025). Cryo-EM structure of the heteromeric TRPC1/TRPC4 channel. Nat Struct Mol Biol, 32(2), 326–338. 10.1038/s41594-024-01408-1

81. Woznica, A., Cantley, A. M., Beemelmanns, C., Freinkman, E., Clardy, J., & King, N. (2016). Bacterial lipids activate, synergize, and inhibit a developmental switch in choanoflagellates. Proceedings of the National Academy of Sciences, 113(28), 7894–7899. doi:10.1073/pnas.1605015113

82. Woznica, A., Gerdt, J. P., Hulett, R. E., Clardy, J., & King, N. (2017). Mating in the Closest Living Relatives of Animals Is Induced by a Bacterial Chondroitinase. Cell, 170(6), 1175–1183.e1111. 10.1016/j.cell.2017.08.005

83. Xu, C., & Min, J. (2011). Structure and function of WD40 domain proteins. Protein Cell, 2(3), 202–214. 10.1007/s13238-011-1018-1

84. Yates, A. D., Allen, J., Amode, R. M., Azov, A. G., Barba, M., Becerra, A., Bhai, J., Campbell, L., Martinez, M. C., Chakiachvili, M., Chougule, K., Christensen, M., Contreras-Moreira, B., Cuzick, A., Fioretto, L. D., Davis, P., De Silva, N. H., Diamantakis, S., Dyer, S., . . . Flicek, P. (2022). Ensembl Genomes 2022: an expanding genome resource for non-vertebrates. Nucleic Acids Research, 50(D1), D996–D1003. 10.1093/nar/gkab1007

85. Zhang, M., Ma, Y., Ye, X., Zhang, N., Pan, L., & Wang, B. (2023). TRP (transient receptor potential) ion channel family: structures, biological functions and therapeutic interventions for diseases. Signal Transduct Target Ther, 8(1), 261. 10.1038/s41392-023-01464-x

86. Zhang, Y., & Skolnick, J. (2005). TM-align: a protein structure alignment algorithm based on the TM-score. Nucleic Acids Research, 33(7), 2302–2309. 10.1093/nar/gki524

87. Zheng, J. (2013). Molecular mechanism of TRP channels. Compr Physiol, 3(1), 221–242. 10.1002/cphy.c120001

