## Supplementary Material for "Diversity and Spatial Segregation of TRP Channels in Choanoflagellates Provide Insight into the Evolutionary Origin of Animal Sensory Systems"

for

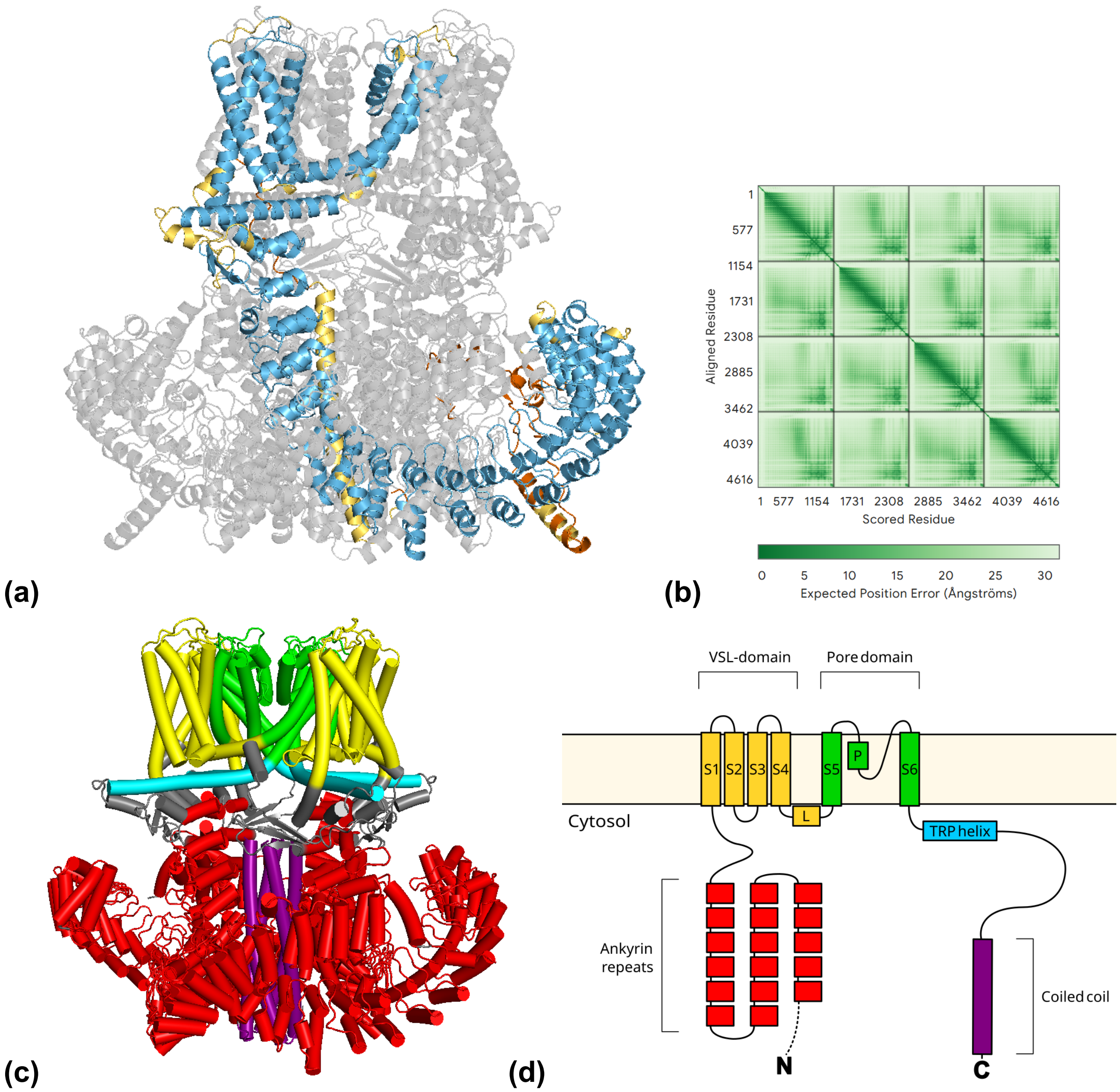


**Figure S1: AlphaFold structure of *sr*TRPA1 (PTSG_04761). (a)** Tetrameric AlphaFold structure with pLDDT values indicated in monomers according to standard coloring scheme (dark blue: >90; light blue: 70-90; yellow: 50-70; orange: <50). **(b)** Predicted alignment error (PAE) for the AlphaFold structure. **(c)** Simplified cartoon representation of *sr*TRPA1 predicted tetrameric structure. Colors correspond to schematic in (d). The region N-terminal to the ankyrin repeats was not included in structure as it was predicted to be intrinsically disordered by InterPro **(d)** Predicted domain topology based on InterPro and AlphaFold structure. Intrinsically disordered regions indicated by a dashed line.

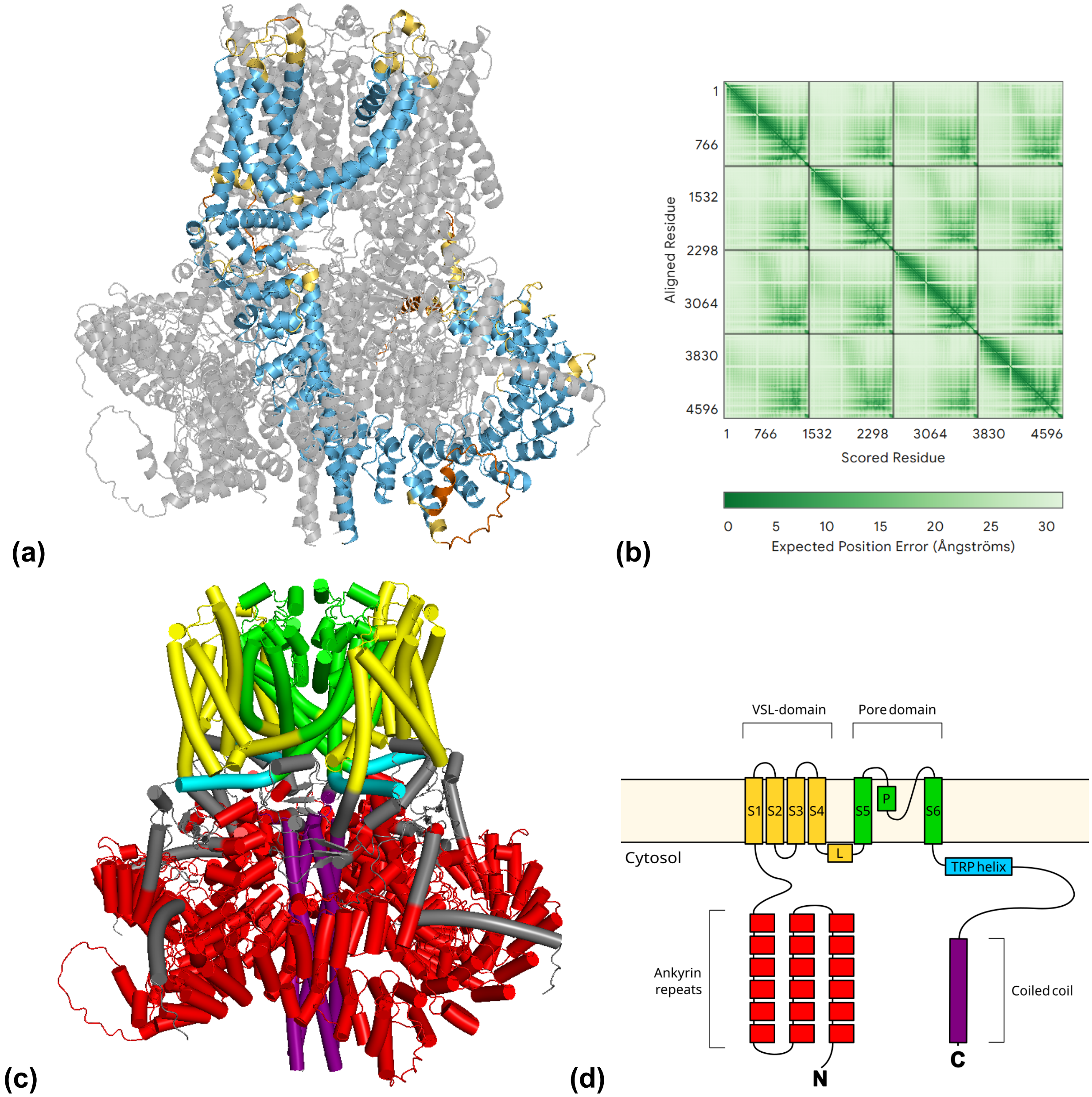


**Figure S2: AlphaFold structure of *sr*TRPA2 (PTSG_08626)**. **(a)** Tetrameric AlphaFold structure with pLDDT values indicated in monomers according to standard coloring scheme (dark blue: >90; light blue: 70-90; yellow: 50-70; orange: <50). **(b)** Predicted alignment error (PAE) for the AlphaFold structure. **(c)** Simplified cartoon representation of *sr*TRPA2 predicted tetrameric structure. Colors correspond to schematic in (d). **(d)** Predicted domain topology based on InterPro and AlphaFold structure.

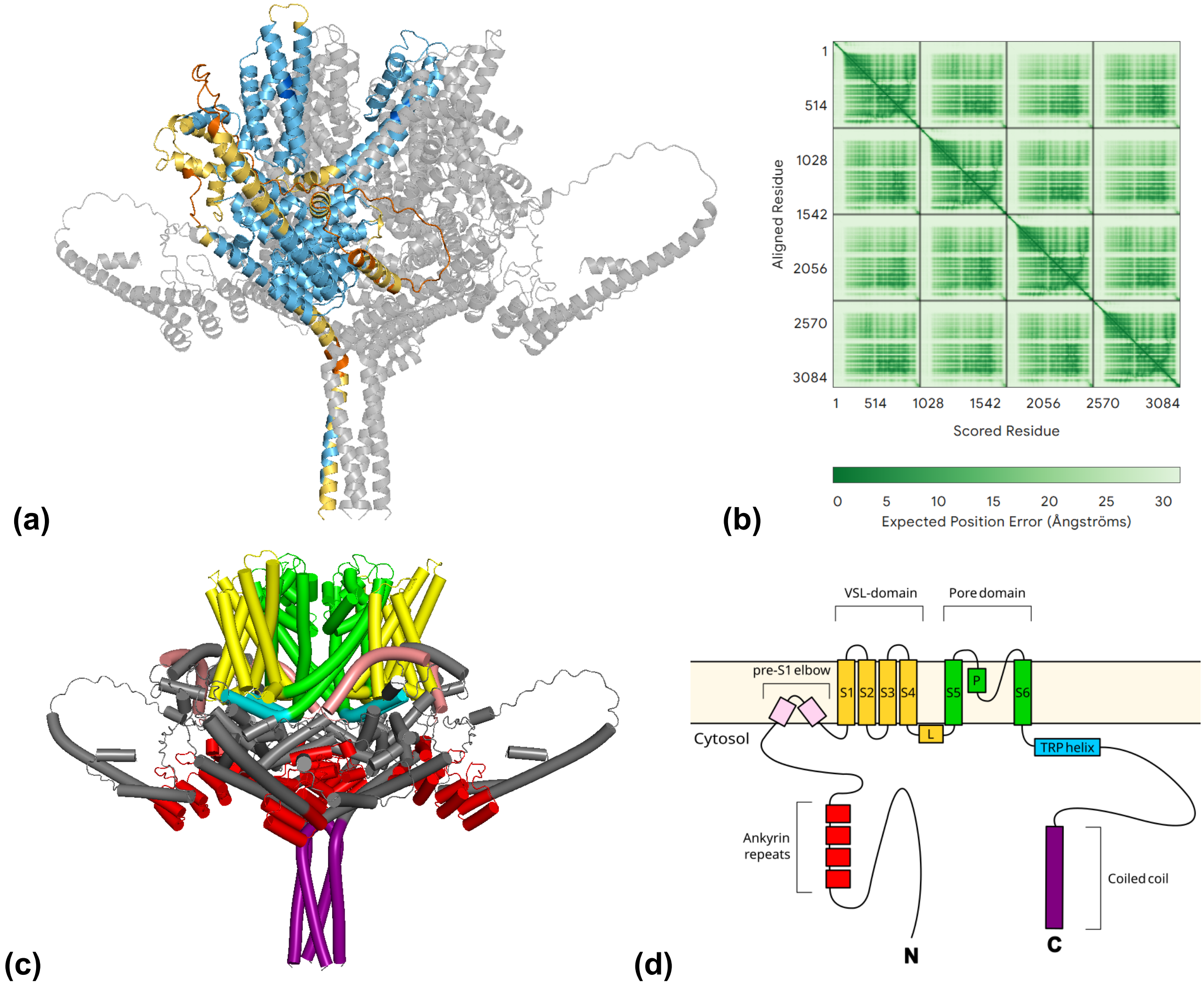


**Figure S3: AlphaFold structure of *sr*TRPC (PTSG_10078)**. **(a)** Tetrameric AlphaFold structure with pLDDT values indicated in monomers according to standard coloring scheme (dark blue: >90; light blue: 70-90; yellow: 50-70; orange: <50). **(b)** Predicted alignment error (PAE) for the AlphaFold structure. **(c)** Simplified cartoon representation of *sr*TRPC predicted tetrameric structure. Colors correspond to schematic in (d). **(d)** Predicted domain topology based on InterPro and AlphaFold structure.

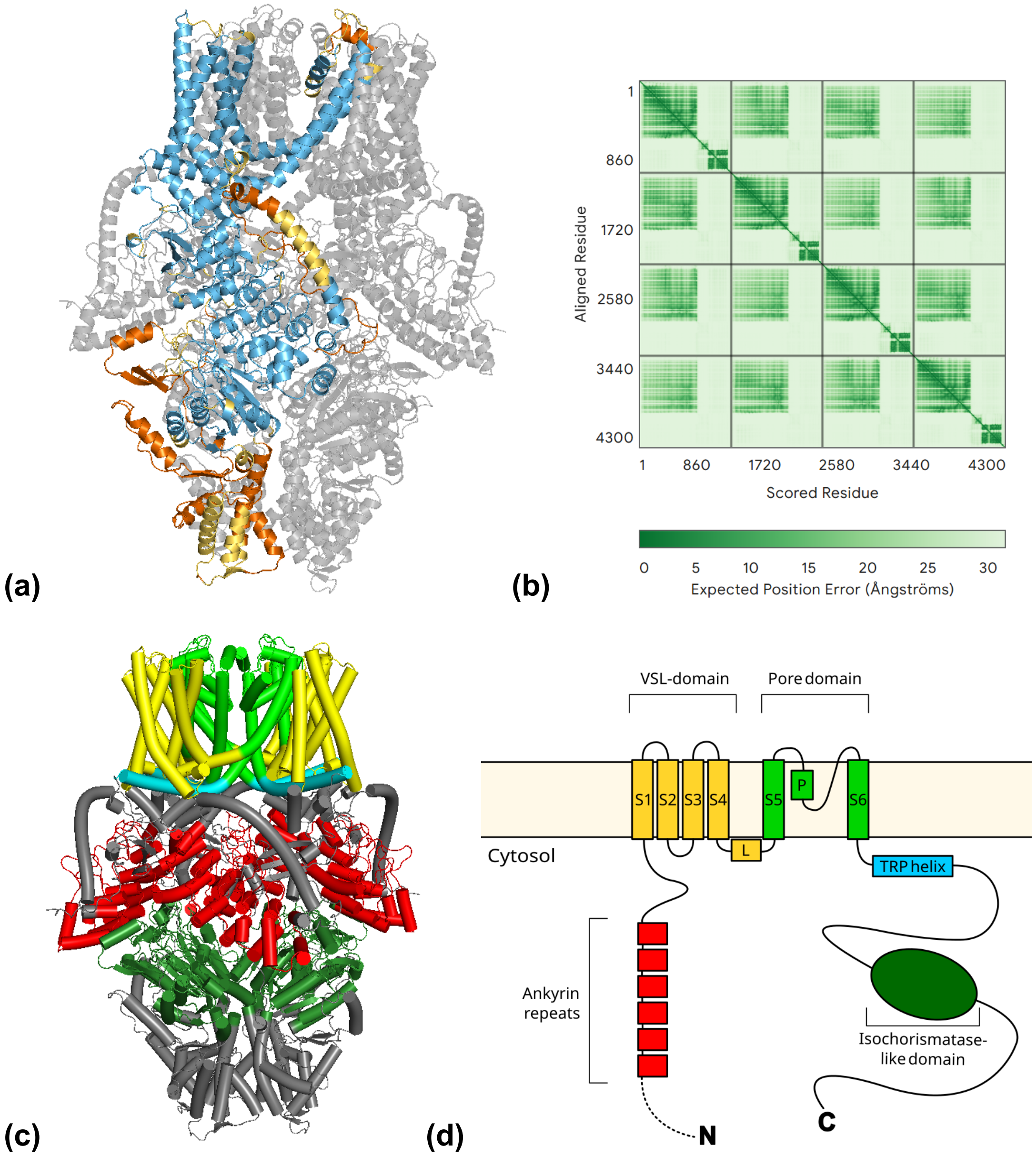


**Figure S4: AlphaFold structure of *sr*TRPV (PTSG_00895). (a)** Tetrameric AlphaFold structure with pLDDT values indicated in monomers according to standard coloring scheme (dark blue: >90; light blue: 70-90; yellow: 50-70; orange: <50). **(b)** Predicted alignment error (PAE) for the AlphaFold structure. **(c)** Simplified cartoon representation of *sr*TRPV predicted tetrameric structure. Colors correspond to schematic in (d). The region N-terminal to the ankyrin repeats was not included in structure as it was predicted to be intrinsically disordered by InterPro **(d)** Predicted domain topology based on InterPro and AlphaFold structure. Intrinsically disordered regions are marked by a dashed line.

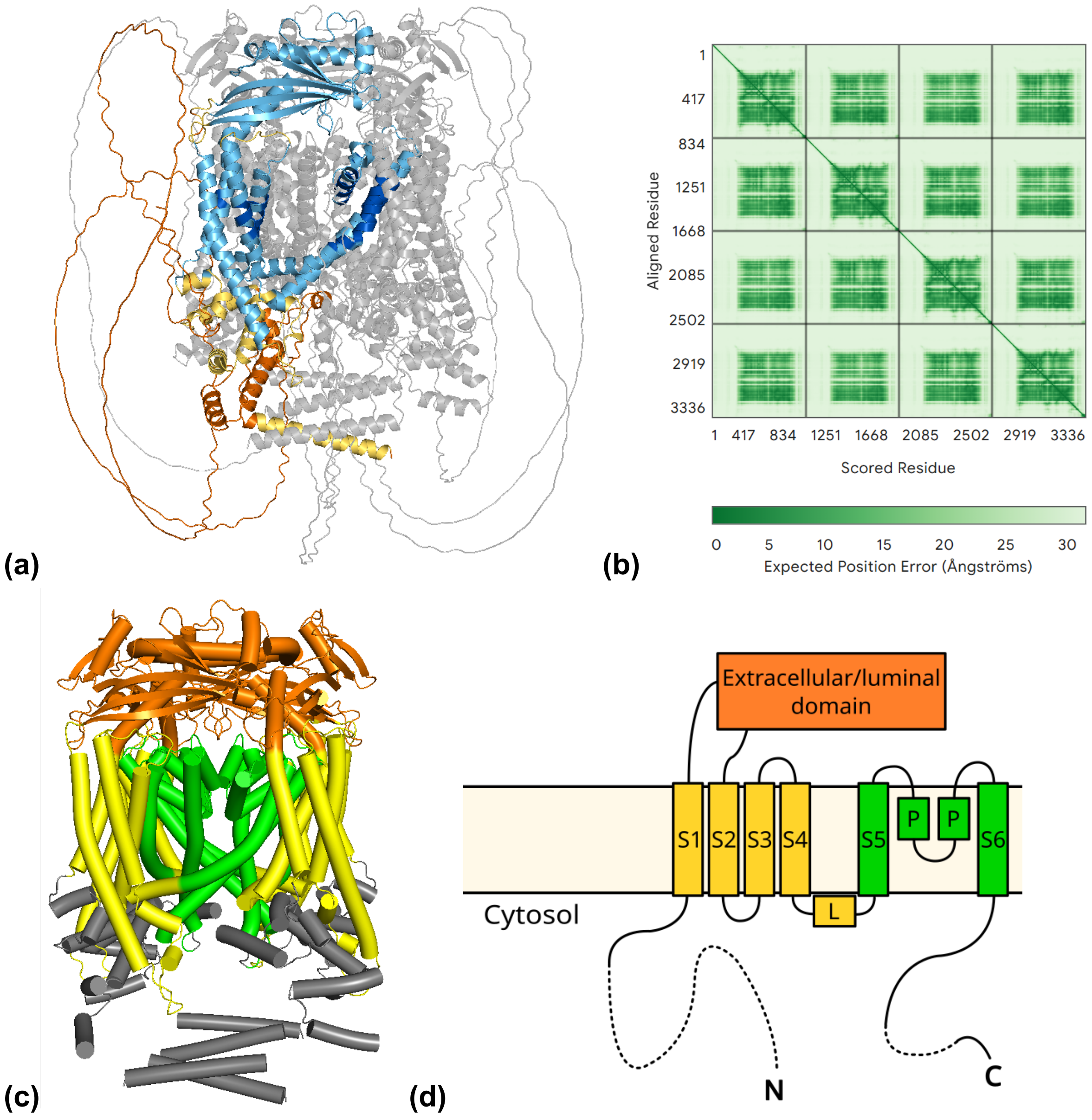


**Figure S5: AlphaFold structure of *sr*TRPML (PTSG_06359)**. **(a)** Tetrameric AlphaFold structure with pLDDT values indicated in monomers according to standard coloring scheme (dark blue: >90; light blue: 70-90; yellow: 50-70; orange: <50). **(b)** Predicted alignment error (PAE) for the AlphaFold structure. **(c)** Simplified cartoon representation of *sr*TRPML predicted tetrameric structure. Colors correspond to schematic in (d). Regions predicted to be intrinsically disordered were not included **(d)** Predicted domain topology based on InterPro and AlphaFold structure. Intrinsically disordered regions are marked by a dashed line.

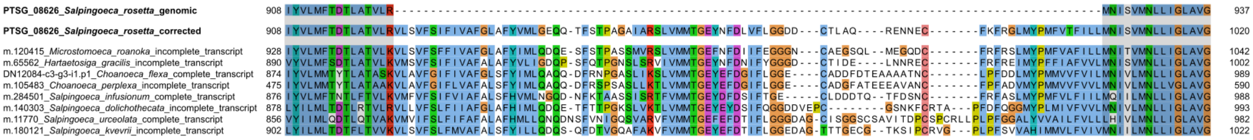


**Figure S6: MSA of S4-S5 linker region for choanoflagellate TRPA2s, including the genomic and the corrected sequence for *sr*TRPA2.** The genomic sequence (PTSG_08626) was predicted to lack the S4-S5 linker region, which would disrupt the TMD. The sequence was corrected by targeted sequencing of cDNA for *sr*TRPA2.

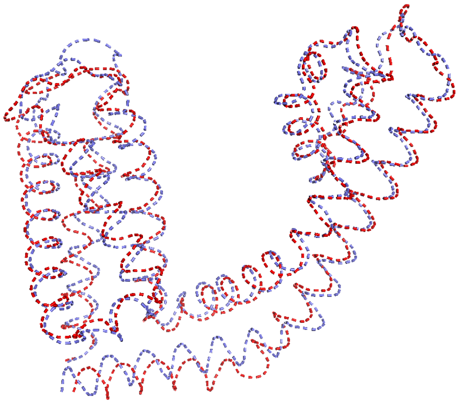

***sr*TRPA1 vs. *hs*TRPA1**
TM score: 0.85
Aligned length: 252 aa
RMSD = 2.23 Å

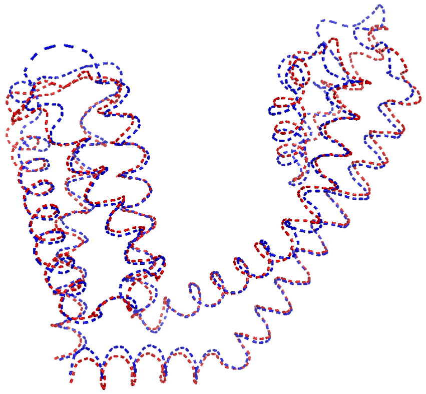

***sr*TRPA2 vs. *hs*TRPA1**
TM score: 0.91
Aligned length: 258 aa
RMSD = 1.70 Å

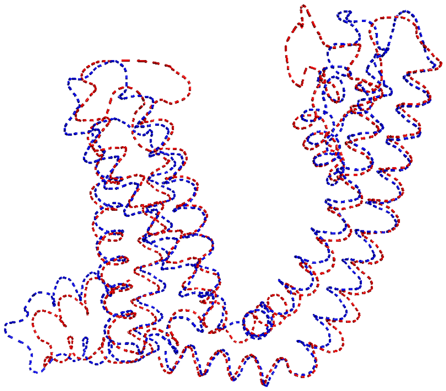

***sr*TRPC vs. *hs*TRPC1**
TM score: 0.79
Aligned length: 289 aa
RMSD = 3.03 Å

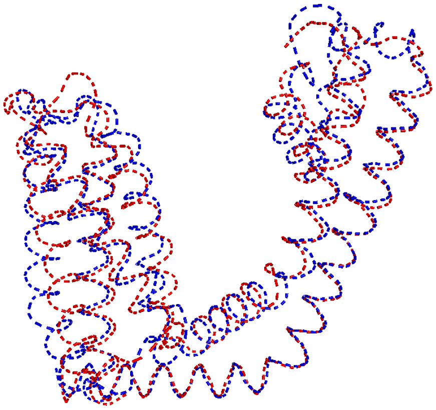

***sr*TRPV vs. *hs*TRPV1**
TM score: 0.88
Aligned length: 251 aa
RMSD = 2.26 Å

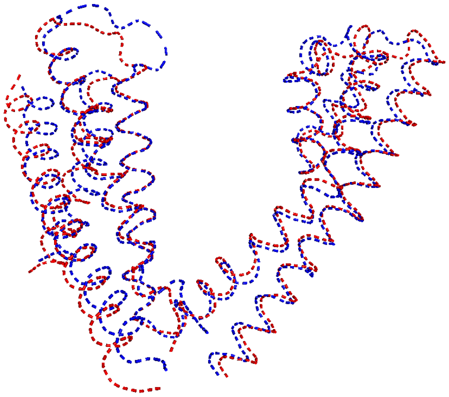

***sr*TRPML vs. *hs*TRPML2**
TM score: 0.89
Aligned length: 246 aa
RMSD = 1.98 Å

**Figure S7 Structural alignments of predicted *S. rosetta* TRP channel TMDs (blue) and experimentally derived TMDs of TRP channels in *Homo sapiens* (red).** TRP helices and pre-S1 elbows were also included in the structural alignment when present. The alignments were done by TMalign, and values were given for the TM scores (on a scale from 0 to 1, and normalized by the structure of *H. sapiens*), number of aligned residues, and the root mean square deviation (RMSD) of aligned residues. The *S. rosetta* structures were predicted by AlphaFold (see **Figure S1-S5**), while the human sequences were downloaded from the protein data bank (RCSB PDB) and derived from cryo-EM data (*hs*TRPA1: 6PQQ; *hs*TRPC1: 9KHJ; *hs*TRPV1: 3J5P; *hs*TRPML2: 9EKW)

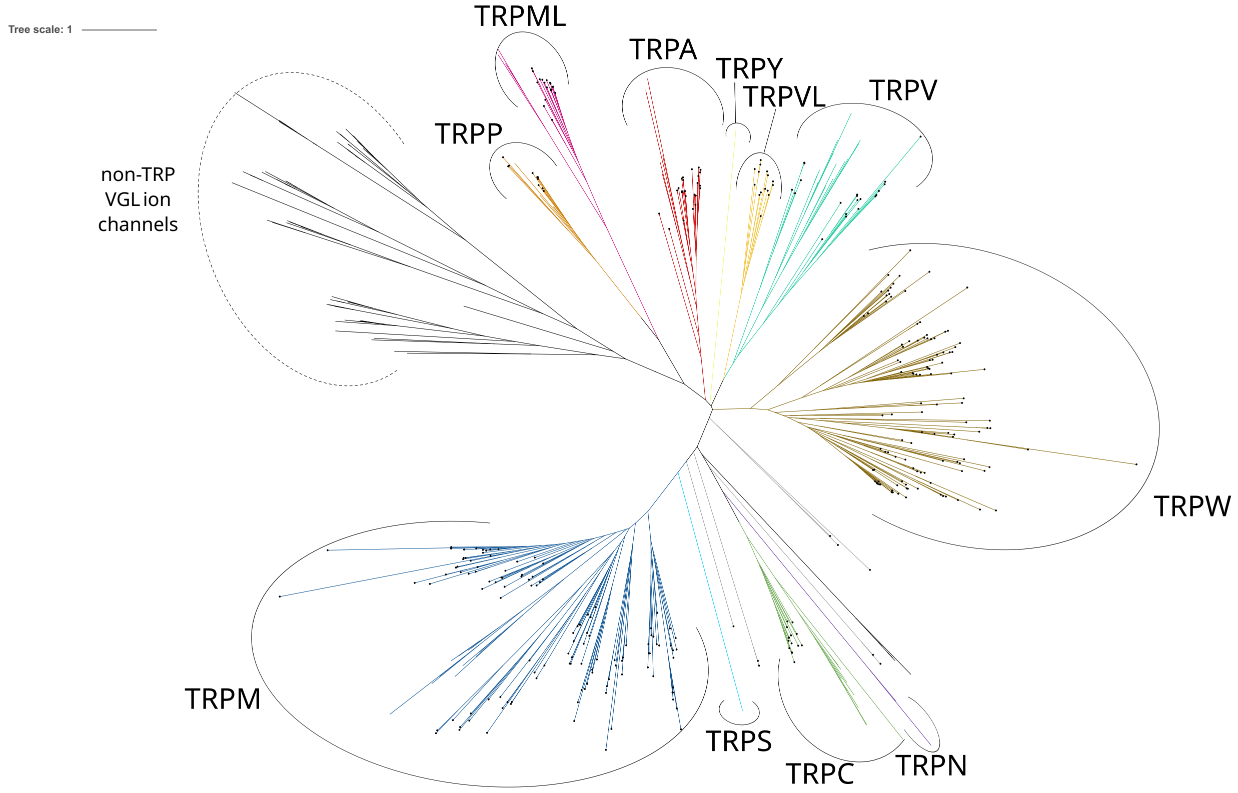


**Figure S8: Phylogeny of choanoflagellate TRP channels.** A maximum-likelihood phylogenetic tree including all identified candidate TRP channels from 21 choanoflagellate species, together with other TRP channels and non-TRP voltage-gated like (VGL) ion channels from other organisms *(Saccharomyces cerevisiae*, *H. sapiens*, *C. elegans* and *D. melanogaster*). The tree was constructed based on the 6TM helix TMD and the 50 flanking residues on each side. All choanoflagellate entries are marked with a black dot on their leaf. The tree with all leaves labelled is available in **Supplemental file S5**.

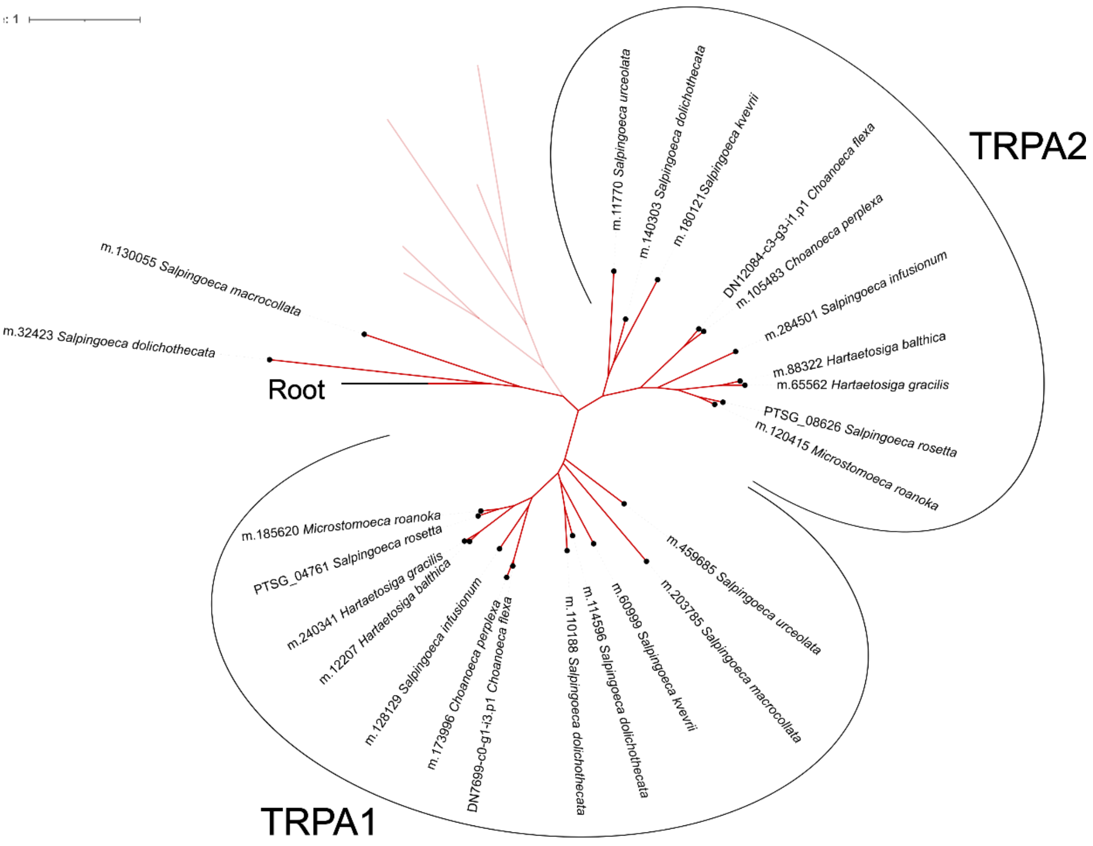


**Figure S9: Phylogeny of the TRPA clade in choanoflagellates**. The shown data was pruned from the TMD-centred maximum-likelihood phylogenetic tree in **Figure S8**. Choanoflagellate entries are highlighted by a black dot at the leaf, while non-marked leaves with opaque branches represent TRPAs from *H. sapiens, D. melanogaster* and *C. elegans* for reference.

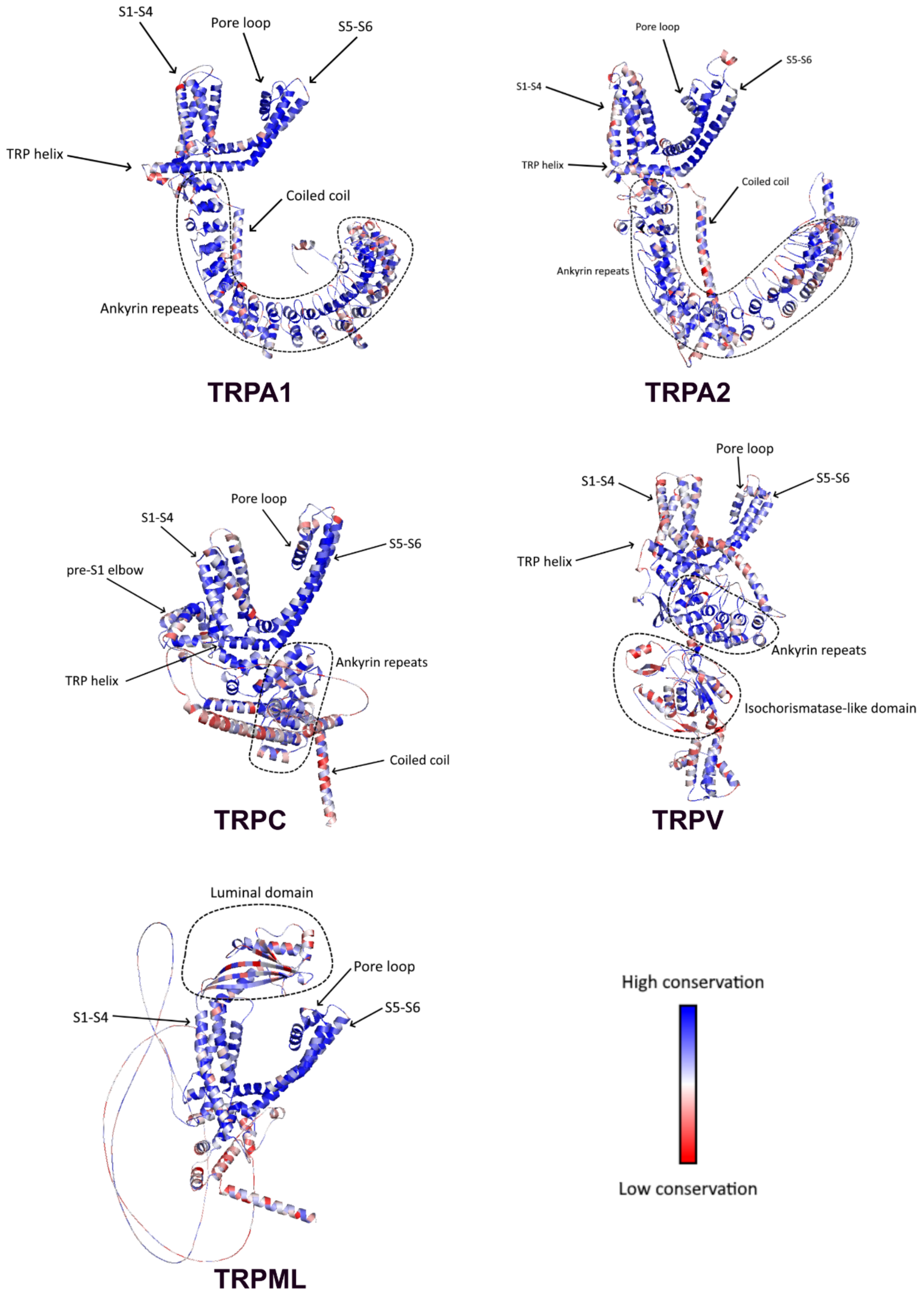
S1-S4
Pore loop
S5-S6
TRP helix
Coiled coil
Ankyrin repeats
TRPA1
Pore loop
S5-S6
S1-S4
TRP helix
Coiled coil
Ankyrin repeats
TRPA2
Pore loop
S1-S4
pre-S1 elbow
S5-S6
TRP helix
Ankyrin repeats
Coiled coil
TRPC
Pore loop
S1-S4
S5-S6
TRP helix
Ankyrin repeats
Isochorismatase-like domain
TRPV
Luminal domain
Pore loop
S1-S4
S5-S6
TRPML
High conservation
Low conservation

**Figure S10: Residue conservation in TRP channel clades in choanoflagellates.** Residue-specific conservation among choanoflagellate TRPA1s, TRPA2s, TRPCs, TRPVs and TRPMLs were mapped onto the predicted structure of *S. rosetta* members, with only one monomer being shown. The full-length protein sequences were aligned, and the degree of conservation was calculated based on the rate4site algorithm. Note that residues in the MSA where *S. rosetta* sequence has gaps cannot be mapped onto the structures and are therefore not shown.

### TRPA1

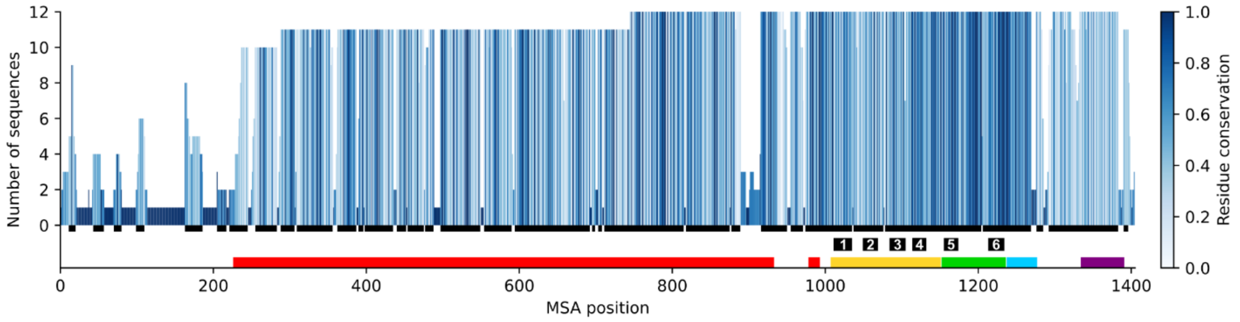


### TRPA2

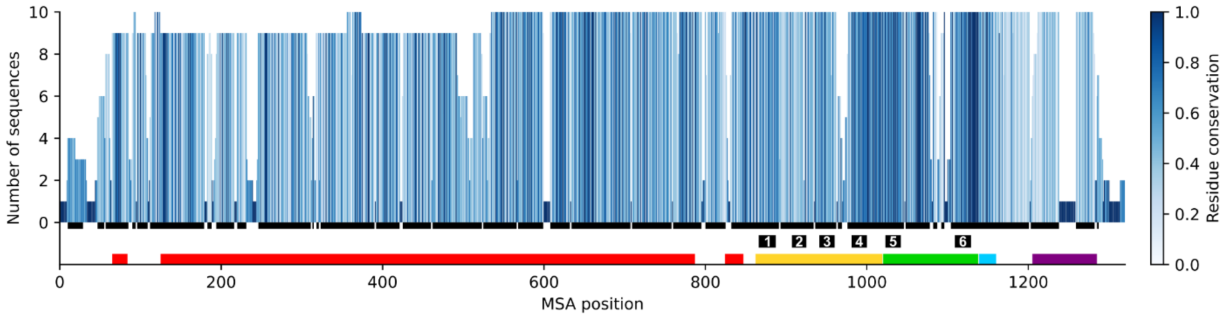


### TRPC

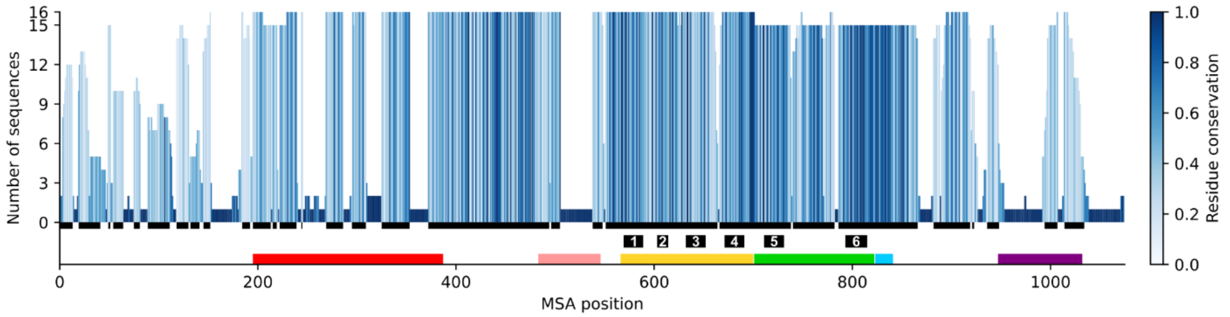


### TRPV

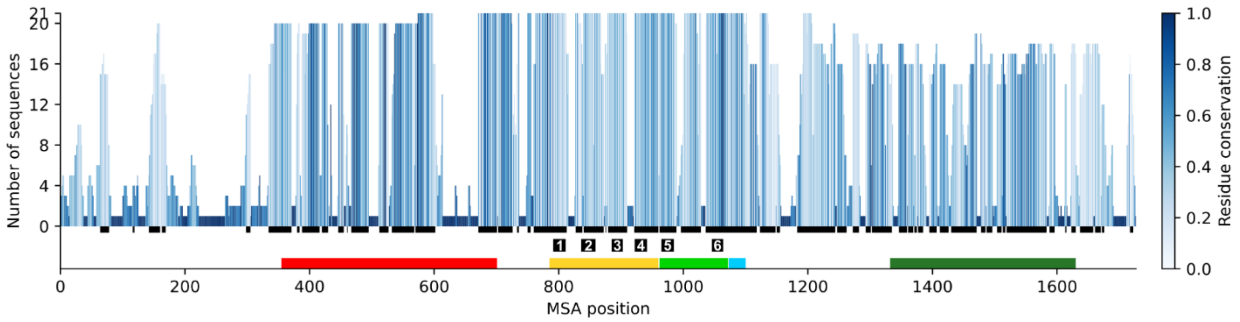

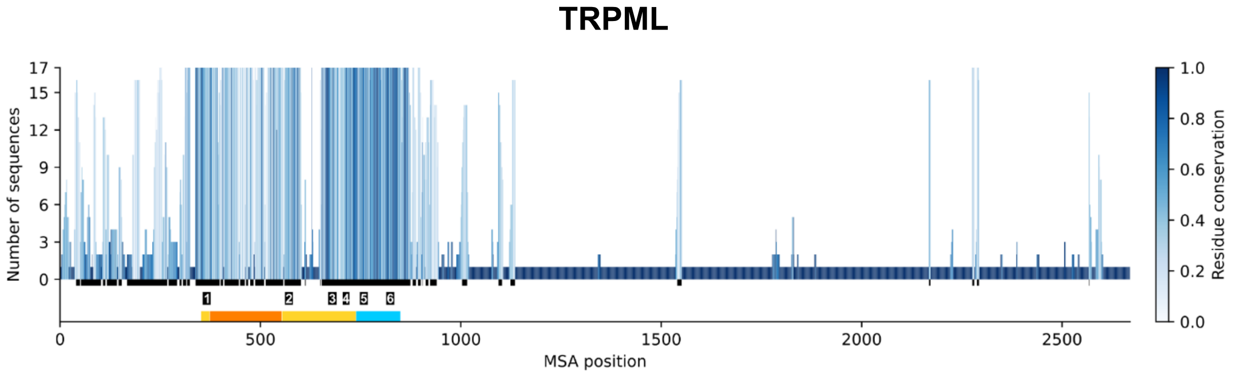


**Figure S11: Summary of clade-specific MSAs of choanoflagellate TRP channels.** To account for the absence of some MSA positions in **Figure S10**, the full MSAs of all choanoflagellate sequences within the clades TRPA1, TRPA2, TRPC, TRPV and TRPML were summarized by the number of sequences represented at each position (bar height) and the residue conservation (color: calculated as normalized, inverted Shannon entropy). The residues included in **Figure S10** were shown by dark grey horizontal bars directly below the graph. The predicted transmembrane TM helices and domain content for the *S. rosetta* entries were also indicated by numbers S(1-6) and colors (corresponding to **Figure S1-S5:** ankyrin repeats: red; pre-S1 elbow: pink; VGL domain: yellow; ion-forming domain: light green; TRP helix: light blue; coiled coil: purple; isochorismatase-like domain: dark green; luminal domain: orange), respectively. Note that some sequences in the MSA are derived from incomplete transcripts.

**Table S1 - List of primers**

| Local Primer ID | Sequence | Application |
|---|---|---|
| **Primers for TRP channel overexpression** | | |
| SM017_pSG_C_F | ATGGCCTCCACCCCCTTCAAG | Amplifying StayGold |
| SM020_pSG_N_R | GAGGTGGGCCTCGAGGGTC | Amplifying StayGold |
| SM021_TRPA1_SGC_F | TCCCAGACCACAACCACAAAACAACCAGCCAT GGCTTCCCGGATGAGCTCTGTTGATCCG | Amplifying *sr*TRPA1 cDNA |
| SM022_TRPA1_SGC_R | GAGCTGGAACTTGAAGGGGGTGGAGGCCATGG AGCCACCGCCGGAGCCACCGCCGTCCCAGTCA AGCTCAATATCATTGGCGTG | Amplifying *sr*TRPA1 cDNA |
| SM023_TRPA2_SGC_F | TCCCAGACCACAACCACAAAACAACCAGCCAT GAACTCGCGCGCGATCACACCGAGGCAC | Amplifying *sr*TRPA2 cDNA |
| SM024_TRPA2_SGC_R | GAGCTGGAACTTGAAGGGGGTGGAGGCCATGG AGCCACCGCCGGAGCCACCGCCAGCGTTGTCA ACATGCTGCTGCAGGCGTTT | Amplifying *sr*TRPA2 cDNA and targeted sequencing of *sr*TRPA2 TMD |
| SM025_TRPC_SGC_F | TCCCAGACCACAACCACAAAACAACCAGCCAT GGCCTTTTTGCAGAGGTGTGCAAGCTTGACG | Amplifying *sr*TRPC cDNA |
| SM026_TRPC_SGC_R | GAGCTGGAACTTGAAGGGGGTGGAGGCCATGG AGCCACCGCCGGAGCCACCGCCCTCATCAGTG AGTGCTGCGATCTTCTGCTC | Amplifying *sr*TRPC cDNA |
| SM027_TRPV_SGC_F | TCCCAGACCACAACCACAAAACAACCAGCCAT GGTGCGCCGCGGCAACAGTGTCACGGAC | Amplifying *sr*TRPV cDNA |
| SM028_TRPV_SGC_R | GAGCTGGAACTTGAAGGGGGTGGAGGCCATGG AGCCACCGCCGGAGCCACCGCCGGCGTGGCTG ATCATGCGAGTGCGACGAAC | Amplifying *sr*TRPV cDNA |
| JJC58F | GAGCTCCCCCCAGCATTATCACG | Amplifying NK802 backbone |
| JJC58R | GGCTGGTTGTTTTGTGGTTGTGG | Amplifying NK802 bakbone |
| JJC59F | CGTGATAATGCTGGGGGGAGCTC | Colony PCR |
| JJC59R | CCACAACCACAAAACAACCAGCC | Colony PCR. Sequencing |
| SG_Linker_F | CCAGTCCGAGACCCTCGAGGCCCACCTCTAAG AGCTCCCCCCAGCATTATCACGTCTACT | Facilitating proper plasmid assembly |
| SG_Linker_R | AGTAGACGTGATAATGCTGGGGGGAGCTCTTA GAGGTGGGCCTCGAGGGTCTCGGACTGG | Facilitating proper plasmid assembly |
| SM022_mCherry_Fwd | ATGGTCTCCAAGGGCGAGGAG | Amplifying mCherry |
| SM023_Linker_mCherry_Rev | CATGTTGTCCTCCTCGCCCTTGGAGACCATGG AGCCACCGCCGGAGCCACCG | Amplifying TRP cDNA and linker from plasmids |
| M13 Reverse primer (N53002) | CAGGAAACAGCTATGAC | Sequencing |

**Table S2 – List of plasmids.** The plasmid sequences were derived from Addgene.

| Plasmid | Reference number | Application |
|---|---|---|
| NK802 | #166056 | Used as backbone for overexpression plasmids. |
| pUC19 | #50005 | Used as carrier plasmid during transfection. |
| NK644 | #109096 | Used to stain inner plasma membrane leaflet. |

**Table S3 – List of synthetic constructs**

| Synthetic construct (ID) | Sequence | Application |
|---|---|---|
| SrStayGold | ATGGCCTCCACCCCCTTCAAGTTCCAGCTCAAGGGCACCATCAAC GGCAAGTCCTTCACCGTCGAGGGCGAGGGCGAGGGCAACTCCCAC GAGGGCTCCCACAAGGGCAAGTACGTCTGCACCTCCGGCAAGCTC CCCATGTCCTGGGCCGCCCTCGGCACCTCCTTCGGCTACGGCATG AAGTACTACACCAAGTACCCCTCCGGCCTCAAGAACTGGTTCCAC GAGGTCATGCCCGAGGGCTTCACCTACGACCGCCACATCCAGTAC AAGGGCGACGGCTCCATCCACGCCAAGCACCAGCACTTCATGAAG AACGGCACCTACCACAACATCGTCGAGTTCACCGGCCAGGACTTC AAGGAGAACTCCCCCGTCCTCACCGGCGACATGAACGTCTCCCTC CCCAACGACGTCCAGCACATCCCCCGCGACGACGGCGTCGAGTGC CCCGTCACCCTCCTCTACCCCCTCCTCTCCGACAAGTCCAAGTGC GTCGAGGCCCACCAGAACACCATCTGCAAGCCCCTCCACAACCAG CCCGCCCCCGACGTCCCCTACCACTGGATCCGCAAGCAGTACACC CAGTCCAAGGACGACACCGAGGAGCGCGACCACATCTGCCAGTCC GAGACCCTCGAGGCCCACCTCTAA | StayGold template for fluorescent overexpression |

**Supplemental file S1: List of reference TRP channels**. These sequences were used for the BLAST search for identifying S. rosetta TRP channels, to create an HMM to target TRP channels in other choanoflagellate species, and used as references in the phylogenetic trees and SSNs. Note that for all except the SSN, the sequences were trimmed to be TMD-focused.

**Supplemental file S2: List of reference non-TRP VGL ion channels**. These sequences were used as references in the phylogenetic trees and SSNs. Note that for phylogenetic trees, the sequences were trimmed to be TMD-focused.

**Supplemental file S3: Overview of the annotations of all candidate choanoflagellate TRP channels.** The information of the first tree columns is derived from the original data: sequence name, species, whether the sequence is complete/incomplete transcript/genomic. The next three columns denote the predicted TRP channel family based on the SSN and phylogenetic tree, respectively, and whether the two give different results. The next seven columns denote whether various domains are predicted to present (denoted by 1/0). The last column lists the predicted number of transmembrane helices, based on deepTMHMM. Note that some regions like the pre-S1 elbow of some TRP channels are in some cases recognized as transmembrane.

**Supplemental file S4: Overview of the annotations of all candidate non-choanozoan holozoan TRP channels.** The information of the first tree columns is derived from the original data: sequence name, species, whether the sequence is complete/incomplete transcript/genomic. The next three columns denote the predicted TRP channel family based on the SSN and phylogenetic tree, respectively, and whether the two give different results. The next seven columns denote whether various domains are predicted to present (denoted by 1/0). The last column lists the predicted number of transmembrane helices, based on deepTMHMM.

**Supplemental file S5:** Detailed phylogenetic tree including all identified choanoflagellate TRP channels.

**Supplemental file S6:** Detailed phylogenetic tree including non-choanozoan holozoan TRP channels.
