## Supplementary figures and images for "Diversity and Spatial Segregation of TRP Channels in Choanoflagellates Provide Insight into the Evolutionary Origin of Animal Sensory Systems"

### Supplemental_file_S5_Choanoflagellate_TRP_phylogeny.pdf

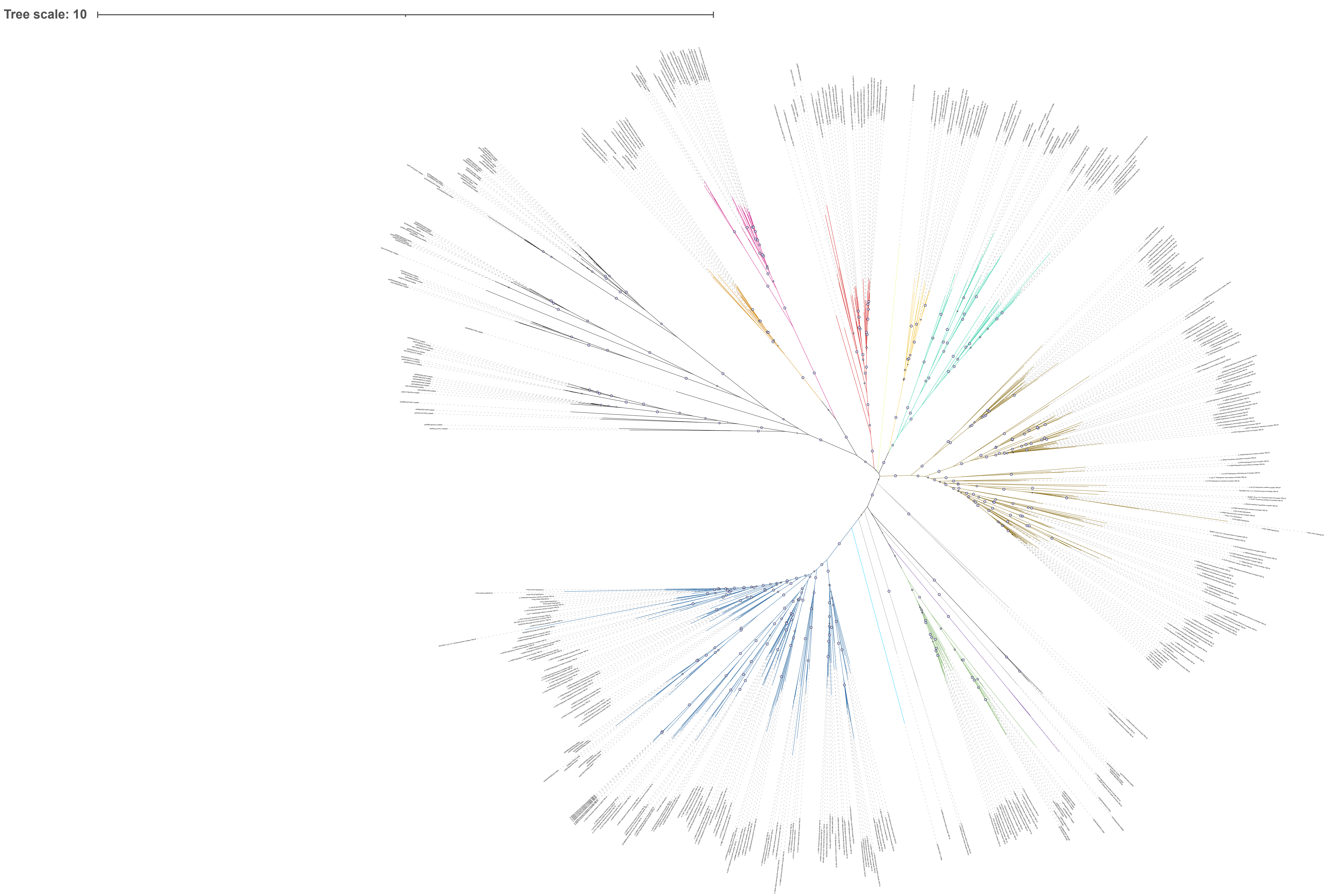

### Supplemental_file_S6_Non-choano_TRP_phylogeny.pdf

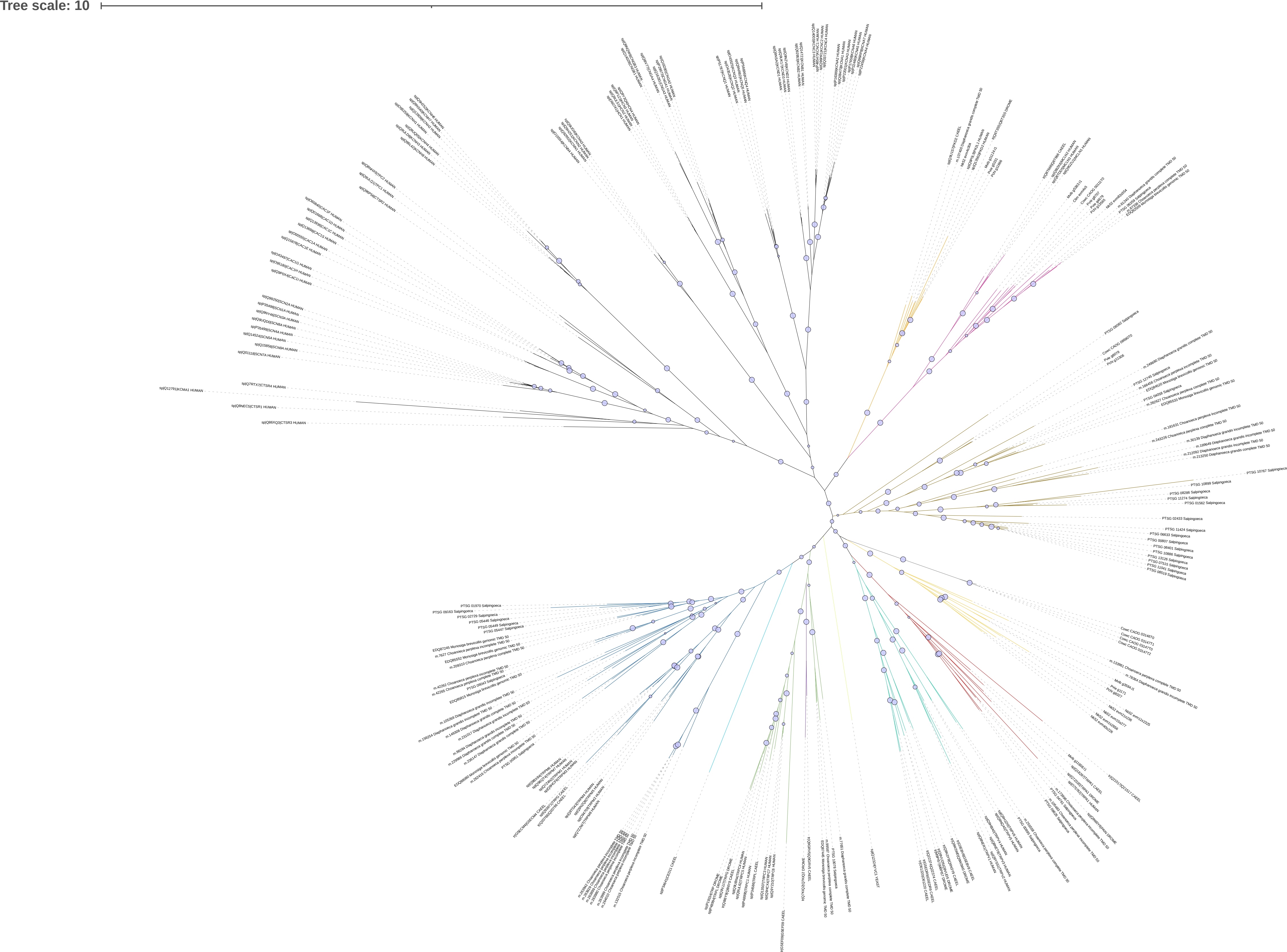
